# Negative allosteric modulation of cannabinoid CB_1_ receptor signaling suppresses opioid-mediated tolerance and withdrawal without blocking opioid antinociception

**DOI:** 10.1101/2024.01.06.574477

**Authors:** Vishakh Iyer, Shahin A. Saberi, Romario Pacheco, Emily Fender Sizemore, Sarah Stockman, Abhijit Kulkarni, Lucas Cantwell, Ganesh A. Thakur, Andrea G. Hohmann

## Abstract

The direct blockade of CB_1_ cannabinoid receptors produces therapeutic effects as well as adverse side-effects that limit their clinical potential. CB_1_ negative allosteric modulators (NAMs) represent an indirect approach to decrease the affinity and/or efficacy of orthosteric cannabinoid ligands or endocannabinoids at CB_1_. We recently reported that GAT358, a CB_1_-NAM, blocked opioid-induced mesocorticolimbic dopamine release and reward via a CB_1_-allosteric mechanism of action. Whether a CB_1_-NAM dampens opioid-mediated therapeutic effects such as analgesia or alters other unwanted side-effects of opioids remain unknown. Here, we characterized the effects of GAT358 on nociceptive behaviors in the presence and absence of morphine. We examined the impact of GAT358 on formalin-evoked pain behavior and Fos protein expression, a marker of neuronal activation, in the lumbar dorsal horn. We also assessed the impact of GAT358 on morphine-induced slowing of colonic transit, tolerance, and withdrawal behaviors. GAT358 attenuated morphine antinociceptive tolerance without blocking acute antinociception. GAT358 also reduced morphine-induced slowing of colonic motility without impacting fecal boli production. GAT358 produced antinociception in the presence and absence of morphine in the formalin model of inflammatory nociception and reduced the number of formalin-evoked Fos protein-like immunoreactive cells in the lumbar spinal dorsal horn. Finally, GAT358 mitigated the somatic signs of naloxone-precipitated, but not spontaneous, opioid withdrawal following chronic morphine dosing in mice. Our results support the therapeutic potential of CB_1_-NAMs as novel drug candidates aimed at preserving opioid-mediated analgesia while preventing their unwanted side-effects. Our studies also uncover previously unrecognized antinociceptive properties associated with an arrestin-biased CB_1_-NAMs.

**Highlights:**

- CB_1_ negative allosteric modulator (NAM) GAT358 attenuated morphine tolerance
- GAT358 reduced morphine-induced slowing of colonic motility but not fecal production
- GAT358 was antinociceptive for formalin pain alone and when combined with morphine
- GAT358 reduced formalin-evoked Fos protein expression in the lumbar spinal cord
- GAT358 mitigated naloxone precipitated withdrawal after chronic morphine dosing

## Introduction

The opioid epidemic remains an issue of great concern that has led to innumerable drug overdose deaths globally[1–3]. While opioids continue to be prescribed for relieving cancer, postoperative and neuropathic pain[4–6], their chronic clinical use is limited by adverse side-effects, including analgesic tolerance, physical dependence, withdrawal, constipation, respiratory depression and, in extreme cases, a fatal overdose[7–12]. Therefore, novel therapeutic strategies that can spare opioid analgesic efficacy while circumventing their unwanted side-effects remain an urgent and unmet therapeutic need. Leveraging the crosstalk between the endocannabinoid and endogenous opioid systems has recently been advanced as one such strategy to achieve this goal [13, 14].

The endocannabinoid system consists of cannabinoid receptors, their endogenous ligands and their respective hydrolytic and synthetic enzymes. This system has been implicated in several opioid-mediated effects including antinociceptive efficacy, tolerance, dependence, withdrawal, and gastrointestinal motility[15–19]. Cannabinoid type 1 receptor (CB_1_) activation via agonists or inhibitors of endocannabinoid deactivation enhances opioid-mediated-antinociception while mitigating tolerance and physical dependence[20–25]. However, the impact of CB_1_ blockade on opioid-mediated therapeutic effects and unwanted side-effects remains incompletely understood. An emerging literature suggests that CB_1_ receptor antagonists or inverse agonists inhibit the development of both acute and chronic opioid analgesic tolerance[26] and naloxone-precipitated opioid withdrawal[27, 28]. Paradoxically, they have also been shown to attenuate mechanical allodynia and thermal hyperalgesia in some preclinical pain models[29–32]. Unfortunately, in patients using rimonabant clinically to treat metabolic syndrome and obesity, unwanted side effects included depression, anxiety, and suicidal ideations, precluding further clinical use [33–36].

The characterization of distinct allosteric binding sites on the CB_1_ receptor[37, 38] has resulted in the development of novel ligands that function as putative CB_1_- positive or negative allosteric modulators (PAMs and NAMs, respectively). The binding of a pure NAM diminishes the affinity and/or efficacy of signaling of the orthosteric agonist or endogenous ligand at the orthosteric binding site, whereas the PAM produces the opposite effect [37, 39–41]. By binding to a site that is distinct from the classical orthosteric binding site, allosteric modulators may theoretically alter endogenous CB_1_ signaling in a spatially and temporally-controlled manner, thereby resulting in a more desirable pharmacological profile overall[42–44]. We recently demonstrated that the CB_1_-NAM GAT358 suppressed opioid-mediated reward at neurochemical and behavioral levels without producing reward or aversion by itself[45]. GAT358 failed to elicit cardinal signs of direct CB_1_ activation or inactivation when administered by itself. Moreover, strikingly, GAT358 did not attenuate cannabinoid CB_1_-mediated antinociception[45]. Thus, CB_1_-NAMs may represent a potential therapeutic strategy for fine-tuning the unwanted pharmacological effects of opioid signaling, while preserving mechanisms required for normal pharmacotherapeutic functions.

In the present study, we used a rat model of inflammatory pain induced by an intraplantar (i.pl.) injection of formalin to evaluate the impact of GAT358 on opioid antinociception. We also evaluated the ability of GAT358 to produce antinociception and suppress neuronal activation in the lumbar spinal dorsal horn using formalin-evoked Fos protein expression as a readout. We also examined the impact of GAT358 in morphine-dependent mice on multiple dependent measures associated with opioid-induced side-effects including slowing of colonic motility, fecal boli production, anti-nociceptive tolerance. Further, we examined the impact of GAT358 on somatic signs of both naloxone-precipitated and spontaneous opioid withdrawal. Our results indicate that GAT358 did not impede the antinociceptive effects of morphine when given in combination.

GAT358 by itself also suppressed both inflammatory nociception and nociceptive processing when administered by itself. Furthermore, GAT358 blocked morphine-induced slowing of colonic motility, and blunted the development of opioid-mediated antinociceptive tolerance. Finally, GAT358 reduced naloxone-precipitated but not spontaneous opioid withdrawal. Our results show that a CB_1_-NAM can block unwanted opioid-induced side-effects without impacting opioid antinociception.

## Materials and Methods

### Animals

Adult male (weighing 25-30g and ∼12-18 weeks old) C57BL/6J (The Jackson Laboratory, Bar Harbor, ME) wildtype (WT) mice and adult male Sprague Dawley (Envigo, Indianapolis, IN) rats (weighing 200-250g and ∼9-12 weeks old) were used in the present study. All animals were maintained on a regular 12h light/dark cycle and given ad libitum access to food (except as noted in the colonic motility experiments) and water. All procedures were approved by the Indiana University Bloomington Animal Care and Use Committee, followed the International Association for the Study of Pain Guidelines for the Use of Animals in Research[46] and complied with the ARRIVE guidelines 2.0[47].

### Drugs

GAT358 (**Supplementary Fig. 1**) and GAT229 were synthesized at Northeastern University and dissolved in a vehicle comprised of dimethyl sulfoxide (DMSO, 20%), ethanol, emulphor, and saline in a 5:2:2:16 ratio, respectively. This vehicle was used to dissolve cannabinoid compounds due to their poor solubility and to assist with drug absorption based on published protocols for in vivo assays using cannabinoid compounds[48]. Previous work from our laboratory has shown that this vehicle did not impact the behavioral parameters assessed in the present study[45, 49–52]. Additionally, the dose of GAT358 (20 mg/kg i.p.) used in the present study was previously shown to reduce morphine-induced dopamine efflux without itself altering dopamine efflux, blocked conditioned place preference to morphine without producing reward or aversion by itself, and failed to elicit the cardinal signs of CB_1_ activation or inactivation [45]. Finally, the dose of GAT229 (20 mg/kg i.p.) used in the present study was previously shown to occlude the behavioral effects of GAT358 in conditioned place preference assays without producing reward or aversion by itself [45]. Morphine sulfate (NIDA Drug Supply Program (Bethesda, MD) or Sigma Aldrich (St. Louis, MO)) was dissolved in saline. All drug and combination treatments were administered via intraperitoneal (i.p.) injection so that the overall injection volumes did not exceed 5 ml/kg in rats and 10 ml/kg in mice, respectively. All test drug pretreatments were performed 20-30 min prior to the morphine injection and control groups received the same vehicle used to dissolve the drug. Formalin was diluted from formaldehyde stock (100% formalin, Sigma Aldrich) in 0.9% saline to a final concentration of 2.5% and administered via an i.pl. injection in a final volume of 50 μL.

### Colonic Motility Bead Expulsion Assay and Fecal Boli Accumulation

The impact of GAT358 on morphine-induced slowing of colonic motility following acute and chronic treatment (on days 1 and 12 of repeated injections, respectively) was measured as described previously by our group [50, 51]. Separate groups of mice received once daily injections of vehicle, GAT358 (20 mg/kg intraperitoneal (i.p.)), morphine (1, 3 or 10 mg/kg i.p.), or the combination of GAT358 (20 mg/kg i.p.) + morphine (1, 3 or 10 mg/kg i.p.) over 12 consecutive days. Mice were fasted for 24 h prior to testing. On the test day, mice were allowed to habituate in the testing room for 30 min prior to testing and were subsequently injected with the test drug. Twenty min following drug administration, a glass bead of approximately 2 mm diameter (Fine Science Tools, Foster City, CA) was inserted 2 cm in the rectal colon using a semi-flexible rubber filament. The time to expel the glass bead (min) was recorded. The weight of fecal boli produced (mg) was also recorded during a 6 h period beginning at the time of the glass bead insertion.

### Hotplate Antinociception

In the same mice used for the colonic assays, the impact of GAT358 on acute antinociception and antinociceptive tolerance to chronic morphine administration (on days 2 and 10 of repeated injections, respectively) was evaluated using the hotplate test as described previously by our group[51, 53]. Separate groups of mice received once daily injections of vehicle, GAT358 (20 mg/kg i.p.), morphine (1, 3 or 10 mg/kg i.p.), or a combination of GAT358 (20 mg/kg i.p.) + morphine (1, 3 or 10 mg/kg i.p.) over 12 consecutive days as described above. A single dose or vehicle condition was evaluated in each group of mice. Hotplate antinociception was measured on day 2 and 10 of repeated dosing to assess both acute drug effects and development of morphine tolerance in a test of acute thermal nociception. Colonic motility (described above) and hotplate antinociception were always measured on different days to minimize stress that could potentially impact assessments. Antinociception was expressed as a percentage of the maximum possible nociceptive effect (% MPE) and was calculated using the following formula: % MPE = [(post morphine latency – baseline latency) / (cut-off latency – baseline latency)] *100[54]. Reduced antinociception on day 10 compared to day 2 of repeated dosing was considered indicative of the development of opioid antinociceptive tolerance.

### Formalin Test

The impact of GAT358 on inflammatory nociception in the formalin test was evaluated as described previously by our group[55, 56]. Separate groups of rats were injected with vehicle, GAT358 (20 mg/kg i.p.), GAT229 (20 mg/kg i.p.), morphine (6 mg/kg i.p.), a combination of GAT358 (20 mg/kg i.p.) + morphine (6 mg/kg i.p.), or a combination of GAT358 (20 mg/kg i.p.) + GAT229 (20 mg/kg i.p.) and placed in an elevated, clear plexiglass observation chamber for a 30 min habituation period. Next, formalin was injected into the superficial plantar surface of either one of the left or right hind paw of all rats via an intraplantar (i.pl.) injection. Formalin-induced nociceptive behaviors were videotaped for 60 mins and subsequently quantified by a trained experimenter who was blinded to the drug conditions. A composite pain score (CPS) was calculated for every 5 min interval during the entire 60 min observation interval using the following scoring criteria: no pain behavior was scored as “0”, lifting of the injected paw was scored as “1”, and shaking/biting/flinching/licking of the injected paw was scored as “2”. The total CPS for each interval was calculated as: [behavior 1 time *+* (2 * behavior 2 time)] / 300. The area under the curve (AUC) of nociceptive behaviors was calculated for the phase 1 (0-10 min) and the phase 2 (10-60 min) for each subject as described previously[55, 56].

### Fos Immunohistochemistry

Immunohistochemical procedures were conducted on tissue obtained from a cohort of rats subjected to the formalin test using procedures described previously by our group[55, 56]. Immediately following the end of the behavioral experiment (∼ 1h post formalin injection), all rats were deeply anesthetized with 25% urethane, and transcardially perfused with 0.1% heparinized 0.1 M phosphate-buffered saline (PBS) followed by ice cold 4% paraformaldehyde (PFA). Lumbar spinal cord tissue was dissected and first placed in PFA for 24 h followed by cryoprotection in 30% sucrose for three days prior to sectioning. Before sectioning, the side of the spinal cord contralateral to the formalin-injected forelimb was marked using an insulin syringe needle to differentiate the hemisections. Transverse sections (40 μm) of the L4 and L5 segments of the lumbar spinal cord were cut on a cryostat and maintained in an antifreeze solution (50% sucrose in ethylene glycol and 0.1 M PBS) prior to immunostaining. Tissue was collected such that every fourth section was processed for immunohistochemistry to prevent the same cell from being inadvertently counted twice in adjacent sections. Free-floating sections were washed in 0.1 M PBS, and subsequently immersed in 0.3% hydrogen peroxide for 30 min. To prevent non-specific binding, sections were pretreated for 1h with a blocking buffer consisting of 5% normal goat serum and 0.3% Triton X-100 in 0.1 M PBS, followed by incubation with rabbit monoclonal anti-c-*fos* antibody (1:10,000; 9F6 #2250 Cell Signaling Technology, Danvers, MA) for 48 h. Sections were washed and incubated in the presence of biotinylated goat anti-rabbit IgG followed by Vectastain elite ABC reagent (1:600, #PK6101, Vector Laboratories, Burlingame, USA). Fos-like immunoreactive cells were visualized with the avidin-biotin peroxidase method using diaminobenzidine as a chromogen. Sections were washed with double-distilled water, slide mounted, air-dried, dehydrated, and cover-slipped with Neo-Mount® (Sigma Aldrich).

### Quantification of Fos immunoreactive cells

Images were captured from the slide-mounted sections using a Leica (Wetzlar, Germany) DM6B microscope and a Leica DFC9000GT digital camera. The specificity of the immunostaining was verified by omission of the primary antibody from the immunostaining protocols. Fos-expressing cells were counted at 5x magnification bilaterally using a computer-assisted image analysis system (LAS AF 2D, Leica). A semi-transparent spinal cord atlas image (**Supplementary Fig. 2**) was superimposed upon the captured image to serve as a reference area and the number of cells were quantified by an experimenter blinded to the treatment condition of the tissue as described previously by our group[53]. The spinal subdivisions subjected to quantification were the superficial dorsal horn (laminae I and II), the nucleus proprius (lamina III and IV), the neck region of the dorsal horn (laminae V and VI), and the ventral horn. Statistical analyses were conducted on the mean number of Fos-positive cells per subdivision averaged across the sections quantified per animal to generate a single mean for each subdivision per animal.

### Naloxone-precipitated Opioid Withdrawal

The impact of GAT358 on naloxone-precipitated opioid withdrawal was evaluated in mice subject to a three-day escalating dosing regimen to induce morphine dependence as described previously by our group[51, 57]. Mice received a total of five i.p. morphine injections across three consecutive days. On days 1 and 2, mice received twice daily (9 am and 5 pm) morphine injections (day 1: 7.5 and 15 mg/kg i.p.; day 2: 30 and 30 mg/kg i.p.). The GAT358+morphine group received a GAT358 injection (20 mg/kg i.p.) 20 min prior to each morphine dose. A separate group received a vehicle injection prior to morphine. On day 3 (48 h following first injection), the body temperature of all mice was measured, and mice were pretreated with either vehicle or GAT358 followed by a final dose of morphine (30 mg/kg i.p.). Next, mice were placed into individual Plexiglas observation chambers to allow for habituation. Then, 30 min (48.5 h) after the final pharmacological treatment, the body temperature was again measured and all mice were challenged with naloxone (10 mg/kg i.p.) to precipitate a µ-opioid receptor-dependent withdrawal syndrome as described previously[51, 57, 58]. Immediately following the naloxone injection, mice were returned to the plexiglass observation cylinders and video recorded continually throughout the observation interval. The number of withdrawal jumps observed over 30 min following the habituation and the naloxone challenge was subsequently quantified by a treatment-blinded rater using the BORIS open source software[59]. Thirty min following the naloxone injection, the body temperature was measured a final time. The impact of precipitated withdrawal on temperature (°C) and body weight (gm) was calculated as a % difference; (body temperature measurement taken ∼30 min following saline/naloxone challenge – pre-challenge body temperature / pre-challenge body temperature)*100. No other withdrawal behaviors were quantified.

### Spontaneous Opioid Withdrawal

The impact of GAT358 on spontaneous opioid withdrawal was evaluated in mice subject to a five-day escalating dosing regimen followed by an activity meter session to quantify potential somatic withdrawal signs as described previously by other groups[55, 56]. Mice received i.p. morphine or saline injections twice daily (9 am and 5 pm) for five consecutive days (day 1: 20 and 20 mg/kg i.p.; day 2: 40 and 40 mg/kg i.p.; day 3: 60 and 60 mg/kg i.p.; day 4: 80 and 80 mg/kg i.p.; day 5: 100 and 100 mg/kg i.p.) as described previously[60, 61]. Separate groups of mice received GAT358 (20 mg/kg i.p.) or vehicle (i.p.) 20 min prior to each morphine/saline dose. Vehicle and saline injections were administered in a volume matched to the active treatments. Twenty-four h after the final injection mice were placed into Omnitech Superflex Node activity meters (Dimensions: 42 x 42 x 30 cm) and were examined for signs of somatic withdrawal using the Fusion 6.5 software (Omnitech Electronics, Columbus, OH) during a 30 min recording interval. The total distance traveled (cm) and the center time (s) (defined as the total time spent in the eight-by-eight square center portion of the activity meter) during the observation interval was recorded. The number of fecal boli accumulated in the activity meter at the end of the 30 min period was recorded. The impact of spontaneous withdrawal on body weight (gm) was calculated as a % difference; (body weight measurement ∼30 min following activity meter test – pre-morphine baseline body weight / pre-morphine baseline body weight)*100. No other withdrawal behaviors were quantified.

### Statistical Analyses

Data were analyzed by one-way, two-way or repeated-measures ANOVA followed by Bonferroni post hoc or by paired or unpaired t-tests or by linear regression analyses, as appropriate. All statistical analyses were performed using GraphPad Prism version 7.05 (GraphPad Software Inc., La Jolla, CA, USA, www.graphpad.com). *p<*0.05 was considered statistically significant.

## Results

### GAT358 impacts chronic but not acute effects of morphine-induced slowing of colonic motility

On day 1 of injections, acute morphine treatment prolonged the glass bead expulsion time in a dose-dependent manner, consistent with morphine-induced slowing of colonic motility, whereas GAT358 treatment failed to alter bead expulsion time overall and the interaction between morphine dose and GAT358 treatment was not significant (Interaction: F_3,40_ = 2.636, *p* = 0.0628, Drug: F_1,40_ = 3.203, *p* = 0.0811, Morphine dose: F_3,40_ = 139.6, *p <* 0.0001; **Fig.1B**). The highest dose of morphine (10 mg/kg i.p.), administered in either the presence or absence of GAT358, was associated with the longest bead expulsion time compared to either vehicle, the low (1 mg/kg i.p.) or middle (3 mg/kg i.p.) dose of morphine (*p <* 0.0001 vs. all other doses). The middle dose of morphine (3mg/kg i.p.), administered in either the presence or absence of GAT358, was also associated with longer bead expulsion time compared to vehicle treatment alone (*p =* 0.0228).

**Fig. 1.**
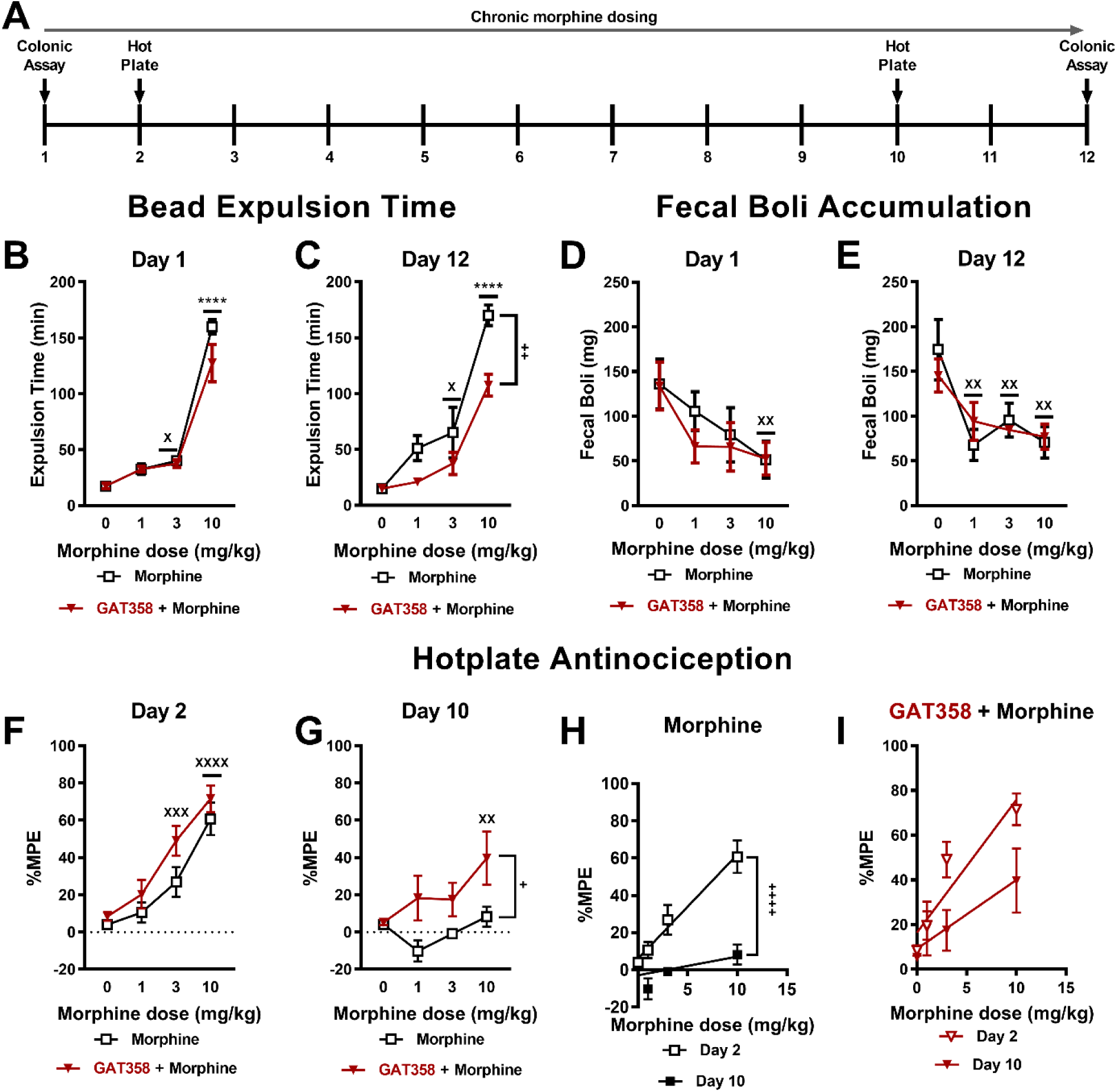
GAT358 attenuates the chronic effects of morphine-induced slowing of colonic motility and development of morphine antinociceptive tolerance in morphine-dependent mice. **(A)** The schematic shows the 12-day chronic dosing paradigm for inducing morphine dependence in mice. Testing days for assessing colonic motility and hot plate antinociception is indicated by black vertical arrows. **(B)** Acute high dose morphine (10 mg/kg i.p. x 1 day) increased bead expulsion times compared to all other morphine dose and vehicle groups. Acute middle dose morphine (3 mg/kg i.p. x 1 day) increased bead expulsion times compared to the vehicle group alone. Bead expulsion times were not affected by GAT358 treatment. **(C)** Chronic high dose morphine (10 mg/kg i.p. x 12 days) increased bead expulsion times compared to all other doses. Chronic middle dose morphine (3 mg/kg i.p. x 12 days) increased bead expulsion times compared to the vehicle group. Bead expulsion times were lower in the GAT358+morphine groups compared to the corresponding morphine alone groups. **(D)** Acute high dose morphine (10 mg/kg i.p. x 1 day) reduced fecal boli accumulation compared to all other doses and was not affected by GAT358 treatment. **(E) High** (10 mg/kg i.p.), middle (3 mg/kg i.p.), and low dose (1 mg/kg i.p.) morphine administered chronically for 12 days reduced fecal boli accumulation compared to vehicle and was not affected by GAT358 treatment. **(F)** Acute high (10 mg/kg i.p. x 2 days) dose morphine, produced higher maximal possible antinociceptive effect (%MPE) in both the morphine alone and GAT358+morphine groups. Acute middle dose morphine (3 mg/kg i.p. x 2 days), produced higher %MPEs in the GAT358+morphine group alone. **(G)** Chronic high dose morphine (10mg/kg i.p. x 10 days) failed to produce antinociceptive effects in the morphine alone group whereas hotplate antinociception was still present in the chronic GAT358+high dose morphine (10mg/kg i.p. x 10 days) group. **(H)** Linear regression analyses of slopes of the antinociceptive responses on day 2 and day 10 of repeated morphine injections alone showed a significant effect of the injection day, indicative of tolerance. **(I)** Linear regression analyses failed to reveal differences in the slopes of the antinociceptive responses on day 2 and day 10 of repeated GAT358+morphine injections. Data are expressed as mean ± S.E.M. (n = 6 per group). “*” indicates main effect of high dose morphine vs. all other groups where **** *p<*0.0001; “X” indicates main effect vs. vehicle treatment alone where XXXX *p<*0.0001, XXX *p<*0.001, XX *p<*0.01 and X *p<*0.05; “*+*” indicates vs. corresponding GAT358+morphine group with the same symbol indications, two-way ANOVA followed by Bonferroni’s post hoc or planned comparisons test.

Twelve days of chronic morphine treatment increased glass bead expulsion time in a dose-dependent manner, consistent with the absence of tolerance to opioid-induced slowing of colonic motility, whereas GAT358 treatment reduced bead expulsion times but the interaction between morphine dose and GAT358 treatment was not significant (Interaction: F_3,37_ = 2.632, *p* = 0.0643, Drug: F_1,37_ = 14.07, *p* = 0.0006, Morphine dose: F_3,37_ = 48.06, *p <* 0.0001; **Fig.1C**). The highest dose of morphine (10mg/kg i.p.), administered in either the presence or absence of GAT358, was associated with the longest bead expulsion time compared to either vehicle, the low (1 mg/kg i.p.) or middle (3 mg/kg i.p.) dose of morphine (*p <* 0.0001 vs. all other groups). The middle dose of morphine (3mg/kg i.p.), administered in either the presence or absence of GAT358, was also associated with longer bead expulsion time compared to vehicle treatment alone (*p =* 0.0148). Notably, the GAT358 + morphine (10 mg/kg i.p.) group showed lower bead expulsion times compared to the corresponding morphine (10 mg/kg i.p.) alone group (*p =* 0.0010).

On day 1 of injections, acute morphine treatment also decreased the fecal boli accumulation in a dose-related manner, consistent with opioid-induced constipation, whereas GAT358 treatment failed to alter fecal boli accumulation and the interaction between morphine dose and GAT358 treatment was not significant (Interaction: F_3,40_ = 0.2825, *p* = 0.8377, Drug: F_1,40_ = 0.6111, *p* = 0.4390, Morphine dose: F_3,40_ = 4.2, *p* = 0.0113; **Fig.1D**). Fecal boli accumulation was lower in the highest dose of morphine (10 mg/kg i.p.), administered in either the presence or absence of GAT358 compared to the vehicle groups (*p =* 0.0090).

Twelve days of repeated morphine treatment reduced fecal boli accumulation in a dose-related manner, consistent with absence of tolerance to opioid-induced constipation, whereas GAT358 treatment failed to alter fecal boli accumulation and the interaction between morphine dose and GAT358 treatment was not significant (Interaction F_3,37_ = 0.6703, *p* = 0.5757, Drug: F_1,37_ = 0.01661, *p* = 0.8982, Morphine dose: F_3,37_ = 7.815, *p* = 0.0004; **Fig.1E**). The high (10 mg/kg i.p.; *p =* 0.0005), middle (3mg/kg i.p.; *p =* 0.0053) and lowest (1mg/kg i.p.; *p =* 0.0024) doses of morphine, administered in either the presence or absence of GAT358, were associated with lower fecal boli accumulation compared to the vehicle groups indicating that GAT358 did not exacerbate or reduce morphine-induced constipation.

### GAT358 does not block the acute antinociceptive effects of morphine but blunts the development of tolerance following chronic morphine dosing in the hotplate test

On day 2 of repeated dosing, both acute morphine (F_3,38_ = 32.95, *p* < 0.0001) and the combination of GAT358+morphine (F_1,38_ = 6.248, *p* = 0.0169) produced antinociception in a dose-dependent manner, but the interaction between the morphine dose and GAT358 dose was not significant (F_3, 38_ = 0.663, *p* = 0.5800; **Fig.1F**). The highest (10mg/kg i.p.) morphine dose increased % maximum possible antinociceptive effect (%MPE) compared to either vehicle (*p* < 0.0001) treatment or the middle (3mg/kg i.p.) (*p* = 0.0042) and low (1mg/kg i.p.) (*p* < 0.0001) doses of morphine. Further, GAT358+morphine (10 mg/kg i.p.) treatment exhibited a greater %MPE compared to the low (1 mg/kg i.p.) dose morphine (*p* < 0.0001) and GAT358 alone (*p* < 0.0001) treatment while the middle (3 mg/kg i.p.) morphine dose exhibited a greater %MPE compared to the low dose morphine (1 mg/kg i.p.) (*p* = 0.0301) and GAT358 alone (*p* = 0.0005) treatment. Moreover, no differences in %MPE were observed for any morphine dose in the morphine alone group compared to the GAT358+morphine groups, indicating that GAT358 treatment did not alter acute opioid antinociception (*p* > 0.05 for all doses).

On day 10 of repeated injections, both morphine (F_3,38_ = 2.902, *p* = 0.0473) and the combination of GAT358+morphine (F_1,38_ = 12.86, *p* = 0.0009) produced dose-dependent antinociception, and the interaction between the morphine dose and GAT358 dose was not significant (F_3,38_ = 1.567, *p* = 0.2132; **Fig.1G**). However, none of the morphine doses increased %MPE relative to vehicle, consistent with the development of tolerance to the antinociceptive effects of morphine. In contrast, the GAT358+morphine (10 mg/kg i.p.) increased %MPE compared to either morphine (*p* = 0.0238) or GAT358 alone (*p* = 0.0173) treatment.

Linear regression analysis revealed that the slope of the dose response curve for producing antinociception was higher on day 2 compared to day 10 of chronic dosing (F_1,43_ = 21.95, *p* <0.0001; **Fig.1H**), consistent with the development of morphine tolerance following repeated dosing. Conversely, the slopes of dosed response curves for producing antinociception did not differ between day 2 and day 10 in the GAT358+morphine groups (F_1,43_ =3.454, *p* = 0.07; **Fig.1I**), consistent with the blunting of tolerance to morphine antinociception by GAT358.

### GAT358 does not impact opioid-induced antinociception in the formalin-induced inflammatory pain assay

An i.pl. injection of formalin increased the CPS of pain behavior in a biphasic manner, drug pretreatments (with morphine alone (i.p.), GAT358 alone (i.p.), or GAT358+morphine (i.p.)) suppressed the CPS overall and the interaction between time and drug treatment was significant (Interaction: F_36,240_ = 4.025, *p*<0.0001, Drug: F_3,20_ = 6.56, *p*=0.0029, Time: F_12,240_ = 8.774, *p* < 0.0001; **Fig. 2A**). Across the observation interval, drug treatments reduced the formalin-evoked CPS from 20-40 min post-injection (*p* < 0.001 for all groups for all timepoints with one exception; GAT358 alone did not suppress CPS at the 40 min post-injection timepoint) compared to vehicle treatment.

**Fig. 2.**
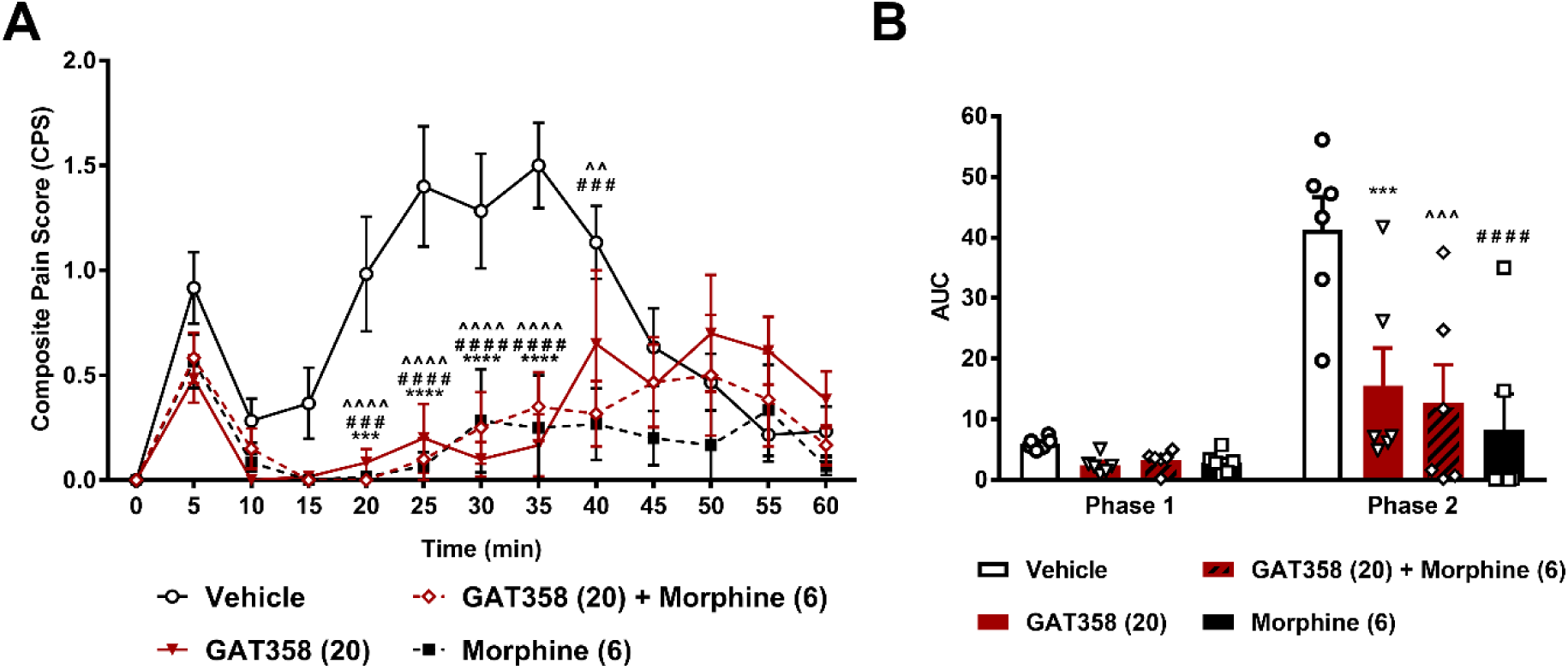
GAT358 does not impact opioid-induced antinociception. **A)** GAT358 (20 mg/kg i.p.) + morphine (6 mg/kg i.p.), morphine alone (6 mg/kg i.p.) or GAT358 alone (20 mg/kg i.p.) suppressed formalin-evoked nociceptive behaviors from 20-40 min following an intraplantar injection of formalin compared to vehicle (i.p.) treatment (with the exception of GAT358 alone at the 40 min timepoint). **B)** All drug treatments reduced the AUC of phase 2 nociceptive behaviors compared to vehicle treatment. Data are expressed as mean ± SEM (n = 6 per group). “*” indicates GAT358 treatment vs. vehicle treatment where **** *p<*0.0001, *** *p<*0.001; “#” indicates morphine treatment vs. vehicle treatment; “^” indicates GAT358+morphine treatment vs. vehicle treatment with the same symbol indications and ^^ *p<*0.01, two-way repeated measures ANOVA followed by Bonferroni’s post hoc test. AUC: area under the curve.

Formalin increased the AUC of nociceptive behaviors in a phase-dependent manner, drug treatments altered the AUC overall, and the interaction between phase and drug treatment was significant (Interaction: F_3,20_ = 5.585, *p* = 0.0060, Drug: F_3,20_ = 6.943, *p* = 0.0022, Phase: F_1,20_ = 31.4, *p* < 0.0001; **Fig. 2B**). All drug treatments reduced the AUC of phase 2 nociceptive behaviors (GAT358+morphine (*p* = 0.0001), morphine (*p* < 0.0001), GAT358 (*p* = 0.0006)), compared to vehicle treatment.

### GAT358 suppresses phase 2 of formalin-evoked nociceptive behaviors and reduced the number of formalin-induced Fos expressing cells in the spinal dorsal and ventral horns

In a separate set of rats subjected to a formalin injection, formalin increased the CPS of pain behavior in a biphasic manner, drug treatments (i.p.) altered the CPS overall and interaction between time and drug treatment was significant (Interaction: F_24,240_ = 3.2, *p* < 0.0001, Drug: F_2,20_ = 13.94, *p* = 0.0002, Time: F_12,240_ = 18.36, *p* < 0.0001; **Fig. 3A**). Across the observation interval, GAT358 treatment reduced the formalin-evoked CPS from 20-35 min post-injection (*p* < 0.01 for all timepoints) compared to vehicle treatment. GAT229 treatment failed to alter the formalin-evoked CPS post-injection compared to vehicle treatment.

**Fig. 3.**
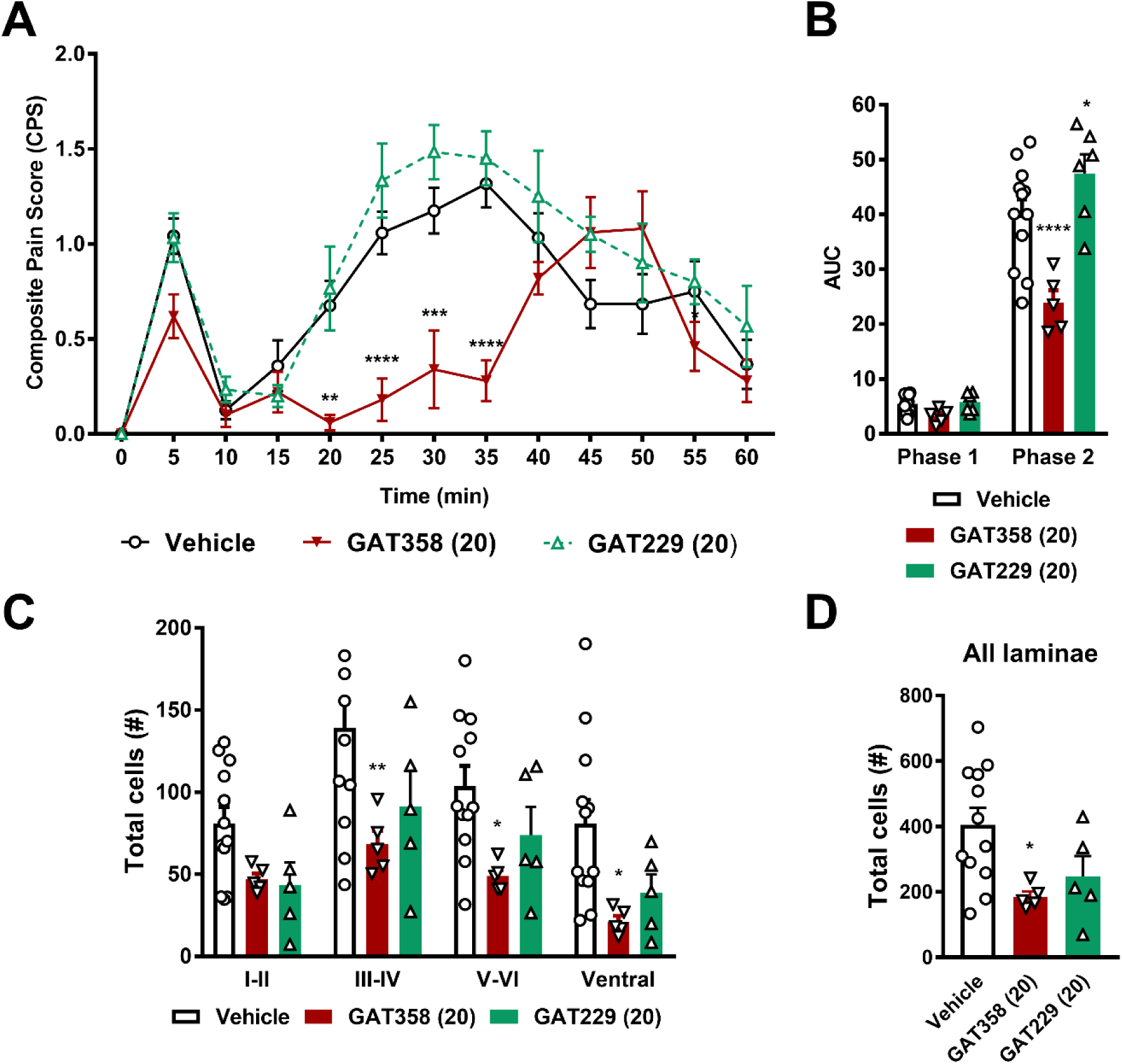
GAT358 but not GAT229 suppresses phase 2 formalin-evoked nociceptive behaviors and reduces the number of formalin-evoked Fos protein expressing cells in the lumbar spinal cord. **A)** GAT358 alone (20 mg/kg i.p.) suppressed formalin-evoked nociceptive behaviors from 20-35 min following an intraplantar injection of formalin compared to vehicle (i.p.) treatment. **B)** GAT358 reduced while GAT229 increased the AUC of phase 2 nociceptive behaviors compared to vehicle treatment. **C)** GAT358 reduced the number of formalin-evoked Fos protein-like immunoreactive cells in the spinal dorsal horn preferentially in laminae III-IV and V-VI and the ventral horn compared to vehicle treatment. **D)** GAT358 reduced the total number of FOS expressing cells in the lumbar spinal cord compared to vehicle treatment. Data are expressed as mean ± SEM (n = 5-12 per group). “*” indicates GAT358 treatment vs. vehicle treatment where **** *p<*0.0001, *** *p<*0.001, ** *p<*0.01, * *p<*0.05, ordinary one-way or two-way repeated measures ANOVA followed by Bonferroni’s post hoc test or planned comparisons test. AUC: area under the curve.

Formalin increased the AUC of phase 1 and phase 2 nociceptive behaviors in a phase-dependent manner, drug treatments altered the AUC overall and the interaction between phase and drug treatment was significant (Interaction: F_2,20_ = 8.061, *p* = 0.0027, Drug: F_2,20_ = 13.75, *p* = 0.0002, Phase: F_1,20_ = 271.2, *p* < 0.0001; **Fig. 3B**). While GAT358 reduced the AUC of phase 2 nociceptive behaviors (*p* < 0.0001), GAT229 increased the AUC of phase 2 nociceptive behaviors (*p* = 0.0399) compared to vehicle treatment.

In a subset of these rats evaluated for formalin-evoked pain behaviors, drug treatments significantly altered the total number of Fos-like immunoreactive cells in the lumbar spinal cord overall (F_2,19_ = 4.403, *p* = 0.0268; **Fig. 3D**) and GAT358 reduced the total number of Fos-like immunoreactive cells in the lumbar spinal cord compared to vehicle treatment (*p* = 0.0276). Further, while drug treatments altered formalin-evoked Fos-like immunoreactive cells overall and Fos expression differed in a laminae-dependent manner, the interaction between drug treatments and spinal cord laminar Fos expression was not significant (Interaction: F_6,76_ = 0.304, *p* = 0.9330, Drug: F_2,76_ = 15.35, *p* < 0.0001, Lamina: F_3,76_ = 6.056, *p* = 0.0009; **Fig. 3C**). Planned comparisons nonetheless confirmed that, across laminae, GAT358 produced robust reductions in the number of formalin-evoked Fos-like immunoreactive cells in the nucleus proprius (*p* = 0.0032) and the neck region (*p* = 0.0263) of the dorsal horn and the ventral horn (*p* = 0.0146) compared to vehicle treatment. GAT229 treatment failed to alter the number of formalin-evoked Fos-like immunoreactive cells compared to vehicle treatment.

### GAT229 occludes the effects of GAT358 in suppressing formalin-evoked nociceptive behaviors and prevents reductions in the number of formalin-induced Fos expressing cells in the spinal dorsal horn

In a separate set of rats subjected to intraplantar formalin and either a combination treatment of GAT358+GAT229 or GAT358 alone, formalin increased the CPS of pain behavior in a biphasic manner, drug treatments altered the CPS overall and the interaction between time and drug treatment was significant (Interaction: F_24,312_ = 4.507, *p* < 0.0001, Drug: F_2,26_ = 10.13, *p* = 0.0006, Time: F_12,312_ = 19.02, *p* < 0.0001; **Fig. 4A**). Across the observation interval, GAT358 treatment reduced the formalin-evoked CPS at 5 and from 20-35 min post-injection (*p* < 0.01 for all timepoints) compared to vehicle treatment. Furthermore, GAT358 treatment reduced the formalin-evoked CPS at 30 min post-injection (*p* = 0.0142) compared to GAT358+GAT229 treatment. Finally, GAT358+GAT229 treatment reduced the formalin-evoked CPS at 35 min post-injection (*p* = 0.0026) compared to vehicle treatment.

**Fig. 4.**
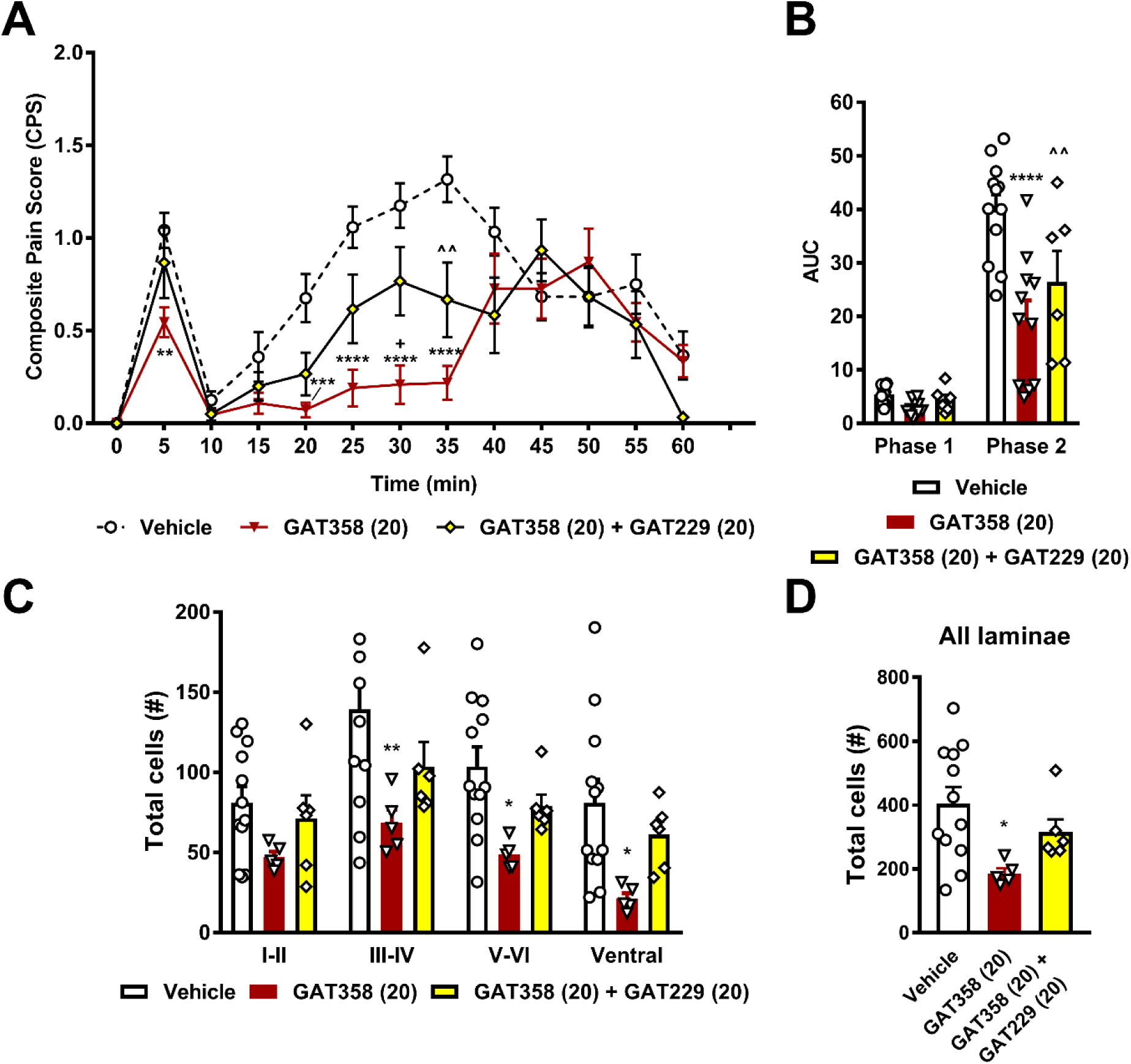
GAT229 occludes the effects of GAT358 on formalin-evoked nociceptive behaviors and formalin-induced Fos expression in the spinal dorsal horn. **A)** GAT358 alone (20 mg/kg i.p.) but not GAT358 (20 mg/kg i.p.) **+** GAT229 (20 mg/kg i.p.) suppressed formalin-evoked nociceptive behaviors following an intraplantar injection of formalin compared to vehicle (i.p.) treatment. **B)** Both GAT358 alone and GAT358**+**GAT229 reduced the AUC of phase 2 nociceptive behaviors compared to vehicle treatment. **C)** GAT358 alone but not GAT358**+**GAT229 reduced the number of formalin-evoked Fos protein-like immunoreactive cells in the spinal dorsal horn preferentially in laminae III-IV and V-VI and the ventral horn compared to vehicle treatment. **D)** GAT358 alone but not GAT358**+**GAT229 reduced the total number of FOS expressing cells in the lumbar spinal cord compared to vehicle treatment. Data are expressed as mean ± SEM (n = 6-12 per group). “*” indicates GAT358 treatment vs. vehicle treatment where **** *p*<0.0001, *** *p*<0.001, ** *p*<0.01, * *p*<0.05; “^” indicates GAT358+GAT229 treatment vs. vehicle treatment; “+” indicates GAT358 treatment vs. GAT358+GAT229 treatment with the same symbol indications, ordinary one-way or two-way repeated measures ANOVA followed by Bonferroni’s post hoc test or planned comparisons test. AUC: area under the curve.

Formalin increased the AUC of phase 1 and phase 2 nociceptive behaviors in a phase-dependent manner, drug treatments altered the AUC overall and the interaction between phase and drug treatment was significant (Interaction: F_2,26_ = 8.633, *p* = 0.0013, Drug: F_2,26_ = 10.41, *p* = 0.0005, Phase: F_1,26_ = 139.4, *p* < 0.0001; **Fig. 4B**). Both GAT358 (*p* < 0.0001) and GAT358+GAT229 (*p* = 0.0031) reduced the AUC of phase 2 nociceptive behaviors compared to vehicle treatment.

In a subset of these rats evaluated for formalin-evoked pain behaviors, drug treatments significantly altered the total number of Fos-like immunoreactive cells in the lumbar spinal cord overall (F_2,20_ = 4.222, *p* = 0.0295; **Fig. 4D**) and GAT358 reduced the total number of Fos-like immunoreactive cells in the lumbar spinal cord compared to vehicle treatment (*p* = 0.0184).

Further, while drug treatments altered formalin-evoked Fos-like immunoreactive cells overall and Fos expression differed in a laminae-dependent manner, the interaction between drug treatments and spinal cord laminar Fos expression was not significant (Interaction: F_6,80_ = 0.3457, *p* = 0.9104, Drug: F_2,80_ = 14.27, *p* < 0.0001, Lamina: F_3,80_= 5.891, *p* = 0.0011; **Fig. 4C**). Planned comparisons nonetheless confirmed that, across laminae, GAT358 produced robust reductions in the number of formalin-evoked Fos-like immunoreactive cells in the nucleus proprius (*p* = 0.0020) and the neck region (*p* = 0.0191) of the dorsal horn and the ventral horn (*p* = 0.0101) compared to vehicle treatment. GAT358+GAT229 treatment failed to alter the number of formalin-evoked Fos-like immunoreactive cells compared to vehicle treatment.

### Representative photomicrographs depict the impact of vehicle and drug treatments on formalin-evoked Fos protein expression in the lumbar spinal cord

Representative photomicrographs depict the impact of vehicle- (**Fig. 5A**), GAT358-treatment (**Fig. 5B**), GAT229-treatment (**Fig. 5C**), and GAT358+GAT229-treatment (**Fig. 5D**) on formalin-evoked Fos protein expression in the lumbar spinal cord ipsilateral to the formalin-injected paw. In the above test rats, there was a relative absence of expression in the dorsal horn of lumbar spinal cord sections contralateral to the formalin-injected paw (data not shown). Furthermore, in control rats, tissue subjected to immunohistochemistry procedures along with all the cohorts described above (**Fig. 3**, **4**), an i.pl. injection of saline in lieu of formalin did not induce appreciable expression of Fos-like immunoreactive cells in the lumbar spinal cord (data not shown), consistent with the results of our previously published studies [55, 62].

**Fig. 5.**
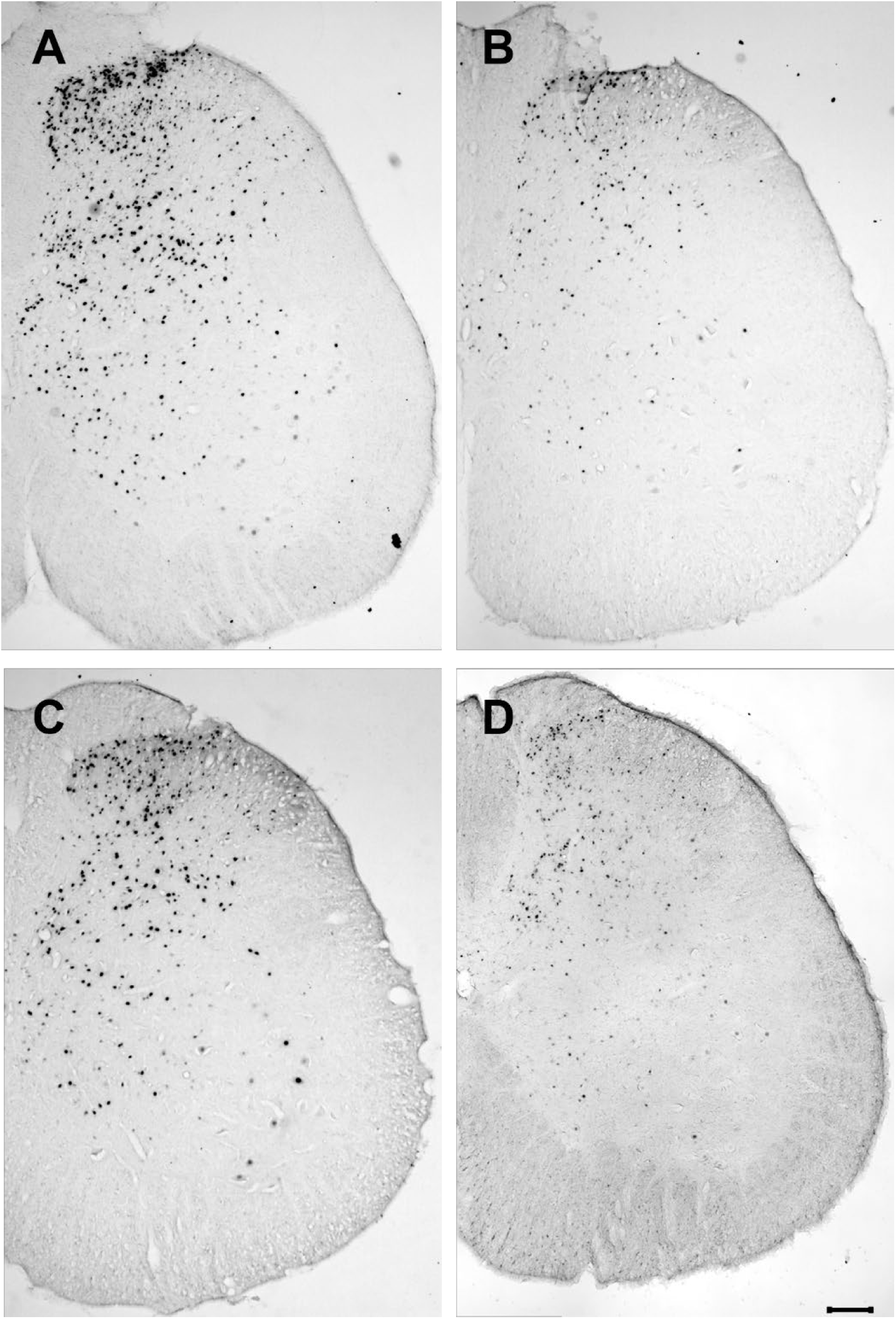
Representative photomicrographs show formalin-evoked Fos immunoreactivity in the lumbar dorsal horn of rats treated with. **A)** Vehicle, **B)** GAT358, **C)** GAT229, **D)** GAT358+GAT229. Scale bar is equal to 100 µm.

### GAT358 treatment in morphine-dependent mice reduces somatic signs of naloxone-precipitated opioid withdrawal

In mice subjected to an escalating i.p. morphine dosing regimen, a naloxone challenge produced characteristic opioid-withdrawal induced jumping behavior compared to the habituation phase, GAT358 (i.p.) reduced the number of withdrawal jumps overall and interaction between challenge and drug treatment was significant (Interaction: F_1,10_ = 9.143, *p* = 0.0128, Drug: F_1,10_ = 9.143, *p* = 0.0128, Challenge: F_1,10_ = 14.17, *p* = 0.0037; **Fig. 6B**). The number of naloxone-precipitated withdrawal jumps was higher in the morphine dependent group that received pretreatment with vehicle (*p =* 0.0014) but not the chronic GAT358-treated group (*p >* 0.05) compared to the same group assessed during the habituation phase (prior to naloxone challenge). Further, the number of naloxone-precipitated jumps was lower in morphine-treated animals that received concurrent treatment with GAT358 (*p =* 0.0007) compared to vehicle (p>0.9999). No differences were observed between the groups following habituation.

**Fig. 6.**
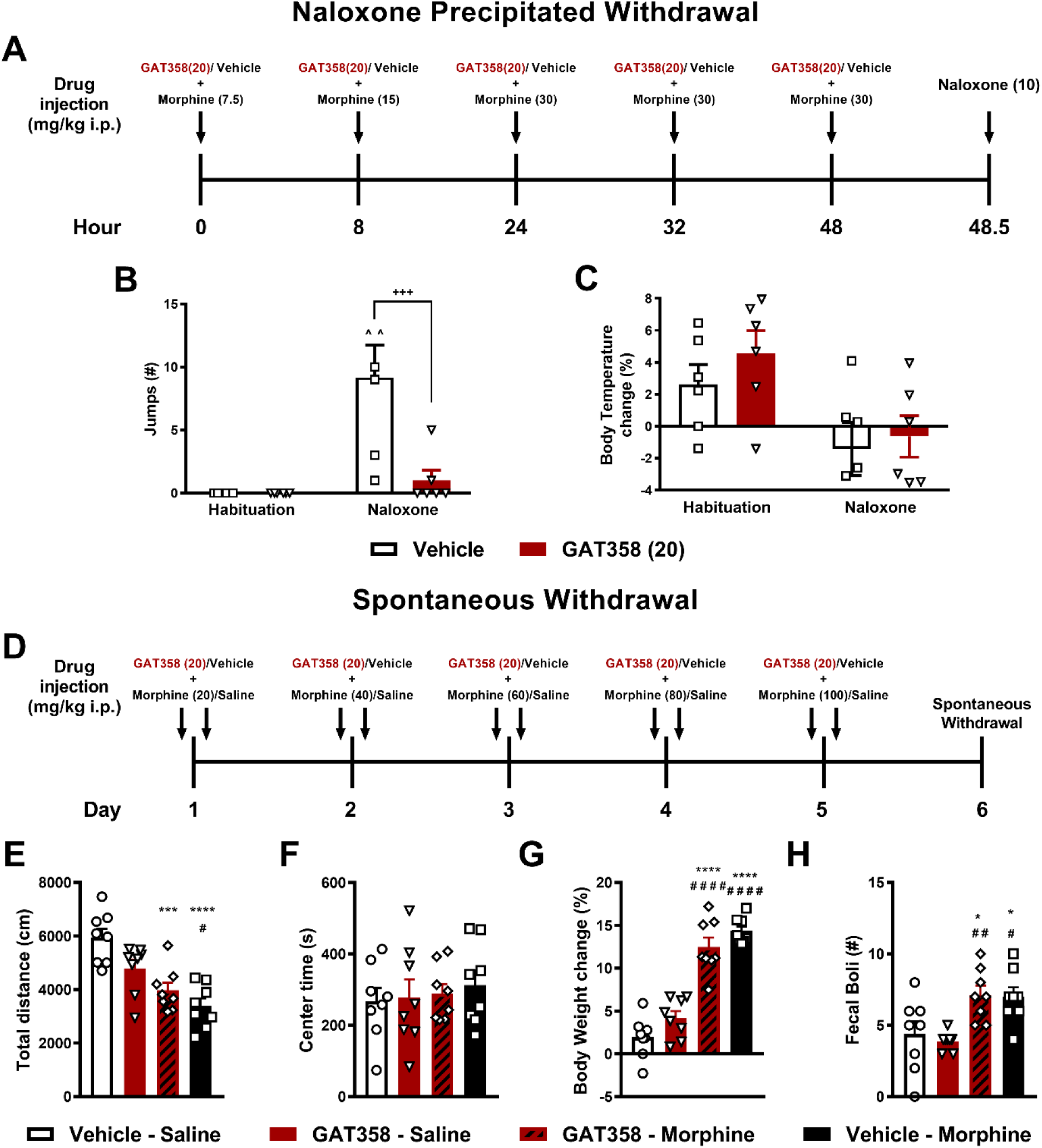
GAT358 attenuates the somatic signs of naloxone precipitated but not spontaneous opioid withdrawal in morphine-dependent mice. **(A)** The schematic shows the timing and doses of the escalating morphine dosing schedule followed by a naloxone (10 mg/kg i.p.) challenge to precipitate opioid withdrawal. **(B)** Naloxone produced characteristic jumping behavior consistent with the development of physical dependence in the vehicle-treated mice. Chronic GAT358-treatment reduced the number of naloxone-precipitated jumps compared to vehicle-treatment. **(C)** Change in body temperature relative to pre-treatment baseline did not differ between groups as a function of challenge condition, indicating absence of body temperature changes following drug treatments or naloxone challenge. **(D)** The schematic shows doses and timing of the escalating morphine dosing schedule followed by behavioral evaluation during a 30 min interval 24 h after the last injection corresponding to spontaneous withdrawal. Morphine-dependent mice showed **(E)** increased locomotor depression but **(F)** no alterations in center time in the activity meter test, **(G)** increased weight loss (%), and **(H)** increased fecal boli accumulation. GAT358 treatment did not alter any of these dependent measures in morphine-treated mice. Data are expressed as mean ± S.E.M. (n = 6-8 per group). *“*”* indicates vs. vehicle-saline where *\*\*\*\*p<*0.0001, *\*\*\*p<*0.001, *\*\*p<*0.01, *\*p<*0.05; “^” indicates vs. habituation; “*+*” indicates vs. corresponding GAT358 treated group; “#” indicates vs. GAT358-saline group with the same symbol indications, ordinary one-way or two-way repeated measures ANOVA followed by Bonferroni’s post hoc test or planned comparisons test.

In these same mice, a naloxone challenge also produced characteristic opioid-withdrawal induced hypothermia compared to the habituation phase, GAT358 (i.p.) failed to alter this withdrawal-induced body temperature change (%) overall and interaction between challenge and drug treatment was not significant (Interaction: F_1,10_= 0.1169, *p* = 0.7395, Drug: F_1,10_= 1.521, *p* = 0.2456, Challenge: F_1,10_= 7.648, *p* = 0.0199; **Fig. 6C**). No reliable differences were found within the groups of mice following either habituation or naloxone challenge or between the chronic GAT358-treated group compared to the vehicle-treated group (*p >* 0.05 for all comparisons).

### GAT358 treatment in morphine-dependent mice does not alter somatic signs of spontaneous withdrawal

In mice subjected to an escalating i.p. morphine dosing regimen, the total distance traveled in the activity meter, assessed 24 h following last injection (i.e., the time period assessed for spontaneous withdrawal) showed a main effect of drug treatment (F_3, 28_ = 12.64, *p* < 0.0001, **Fig.6E**). Vehicle-morphine-treated mice traveled lower total distance compared to GAT358-saline- (*p* = 0.0201) and vehicle-saline-treated (*p* < 0.0001) mice. Further, GAT358-morphine-treated mice traveled a lower total distance compared to vehicle-saline-treated mice (*p* = 0.0007). GAT358-saline treated mice also did not differ from vehicle-saline mice on this measure (*p* > 0.05) indicating the lack of impact of chronic GAT358 treatment on distance traveled in the presence or absence of morphine.

In these same mice, drug treatment did not alter center time in the activity meter (F_3, 28_ = 0.2409, *p* = 0.8670, **Fig.6F**), indicating that this measure was not impacted by discontinuation of morphine treatment or by chronic GAT358 treatment.

In these same mice, body weight change (%) showed a main effect of drug treatment (F_3, 28_ = 51.02, *p* < 0.0001, **Fig.6G**). Vehicle-morphine-treated mice lost more body weight (%) compared to GAT358-saline- (*p* < 0.0001) or vehicle-saline-treated (*p* < 0.0001) mice. Further, GAT358-morphine also lost more body weight (%) compared to GAT358-saline- (*p* < 0.0001) and vehicle-saline-treated (*p* < 0.0001) mice. However, GAT358-saline treated mice did not differ from vehicle-saline mice on this measure (*p* > 0.05), consistent with the lack of impact of chronic GAT358 treatment on this measure in the presence or absence of morphine.

Finally, in these same mice, fecal boli accumulation differed as a function of drug treatment (F_3, 28_ = 6.841, *p* = 0.0013, **Fig.6H**). Vehicle-morphine treated mice produced more fecal boli compared to GAT358-saline- (*p* = 0.0128) and vehicle-saline-treated (*p* = 0.0498) mice. Further, GAT358-morphine groups produced more fecal boli compared to GAT358-saline- (*p* = 0.0090) and vehicle-saline-treated (*p* < 0.0357) mice. However, GAT358-saline treated mice did not differ from vehicle-saline mice on this measure (*p* > 0.05), consistent with the lack of impact of chronic GAT358 treatment on this measure in the presence or absence of morphine.

## Discussion

The Division of Therapeutics and Medical Consequences of the National Institute on Drug Abuse (NIDA) has designated CB_1_-NAMs as one of the top ten most wanted pharmacological mechanisms for the rapid development of therapeutics in response to the opioid crisis[63]. GAT358, a 3,4-diaminocyclobut-3-ene-1,2-dione derivative which shows minimal CB_1_ inverse agonist-related side-effects and displays functional selectivity in the β-arrestin assay[64, 65] was developed by a focused structure-activity relationship study on the classical CB_1_-NAM, PSNCBAM-1[64]. We recently showed that GAT358 blocked morphine-induced reward in a conditioned place preference assay via an allosteric- and CB_1_-mediated mechanism, eliminated morphine-induced evoked dopamine release in the mesocorticolimbic reward pathway and also reduced oral oxycodone consumption without producing observable unwanted side-effects[45]. However, to our knowledge, no prior study has evaluated the in vivo efficacy of a CB_1_-biased NAM on opioid-mediated antinociception, tolerance, and withdrawal. Findings from the current study suggest that negative allosteric modulation of CB_1_ signaling is a potential pharmacological strategy for attenuating opioid-induced tolerance, slowing of colonic motility and naloxone-precipitated opioid withdrawal without impacting its therapeutic antinociceptive effects.

Opioid-induced slowing of colonic motility is a prevalent and debilitating side-effect of chronic opioid use[66, 67], that is reportedly resistant to tolerance despite long-term use[68]. In our study, acute GAT358 treatment failed to alter the inhibitory effects of acute morphine dosing on colonic motility or fecal boli production. However, in a chronic dosing paradigm, GAT358 reduced opioid-induced slowing of colonic motility but not fecal boli accumulation when co-administered with the highest morphine dose. Cannabinoids depress gastrointestinal transit through CB_1_ activation by inhibiting ongoing contractile transmitter release[69, 70]. Reduction of monoacylglycerol lipase either pharmacologically or genetically is shown to accelerate colonic propulsion and makes mice hypersensitive to µ-opioid receptor agonist-mediated inhibition of colonic motility[18]. While rimonabant increased gut motility, small intestinal transit, and caused diarrhea in rodents[69, 71, 72], it failed to block the inhibitory effect of morphine and loperamide on small and whole intestinal transit[73]. More work is necessary to determine the exact mechanisms involved in the chronic but not acute effects of GAT358 treatment on opioid-induced slowing of colonic transit.

Chronic GAT358 treatment blunted development of tolerance to the antinociceptive effects of morphine without altering acute opioid-mediated antinociception. The suppression of opioid-induced increases in calcitonin gene-related peptide implicated in the development of opioid tolerance via spinal CB_1_ receptor blockade may potentially account for this phenomenon[26]. This attenuation of opioid tolerance may prove critical in a clinical setting, as profound individual differences regulate development of tolerance in patient populations, prompting the need for diverse treatment options[74]. Furthermore, the impact of GAT358 is unlike that of direct CB_1_ antagonists and/or inverse agonists such as rimonabant and AM251 which attenuate tolerance but also concomitantly decrease morphine analgesia, thereby decreasing their translational relevance[75, 76].

We also established that GAT358 itself produced antinociception in a test of formalin-induced inflammatory pain and suppressed neurochemical markers of formalin-evoked neuronal activation in regions of the lumbar spinal cord implicated in nociceptive processing[77]. Formalin-evoked nociceptive behaviors occur in a biphasic manner, first with an immediate nocifensive phase 1 beginning 0-5 min post-formalin injection indicative of primary afferent activation[78], and a subsequent phase 2 beginning 15-20 min post-formalin injection, linked to sensitization occurring within the central nervous system and the facilitation of responses to otherwise innocuous stimuli[79, 80]. The time period in between these phases is usually characterized by a state of quiescence and the lack of nociceptive behaviors that may be attributed to transient reductions in primary afferent activity[78] and increased descending inhibitory controls[81, 82]. GAT358 reduced formalin-evoked nocifensive behaviors during phase 1 and phase 2 and also reduced the total number of cells expressing formalin-evoked Fos protein expression, a marker of neuronal activation, in the lumbar spinal cord. This GAT358-induced reduction of Fos was observed both in the nucleus proprius and the neck region of the dorsal horn, that show neuronal activation in response to noxious stimuli[83]. Interestingly, in the spinal cord CB_1_ is also expressed, albeit at low levels, on primary afferents as well as dorsal horn interneurons[30, 84–87]. Notably, the GAT358-induced suppression of Fos expression was observed in the same subjects that exhibited GAT358-induced suppression of formalin-induced pain behavior. These observations support our hypothesis that suppression of formalin-evoked Fos protein expression induced by systemic doses of GAT358 reflect a suppression of nociceptive processing, possibly through enhancing active inhibition during the interphase and/or by inhibiting the contribution of endocannabinoids to activity-dependent pain sensitization[30]. Our results align with a previous report that a CB_1_ antagonist suppressed formalin-induced pain behavior in rats[31]. Moreover, the pattern of changes in pain behaviors and Fos protein expression is similar to that observed previously by our group with other analgesics, including orthosteric cannabinoid agonists[55, 56]. An interesting observation was that the CB_1_ PAM GAT229 did not itself produce antinociception. Intraplantar formalin has been linked to mobilization of the endocannabinoid anandamide in the periaqueductal gray, albeit at a higher concentration (4%) and volume (150 µL) of formalin than that used in the present study[88]. Because activity of PAMs require the presence of the endogenous ligand, it is possible that the pretreatment interval or the concentration and volume of formalin used in the present study were insufficient to reveal a CB_1_ PAM mediated antinociceptive effect.

Because GAT358 was administered systemically in our experiment, our studies do not establish a site of action for the suppression of formalin-evoked neuronal activation and pain behavior. Thus, changes in spinal cord neuronal activation observed here could be either direct or indirect. Several potential mechanisms could mediate this suppression of formalin-evoked pain and spinal Fos protein expression induced by allosteric modulator-mediated disruption of CB_1_ signaling. GAT358 could preferentially suppress endocannabinoid-mediated signaling at GABAergic receptors, where CB_1_ are more densely expressed compared to glutamatergic neurons[87]. GAT358-induced CB_1_ blockade could result in antinociception via non-CB_1_ mediated mechanisms, including shunting of endocannabinoid signaling from CB_1_ to CB_2_ and/or anandamide mediated desensitization of transient receptor potential vanilloid type 1 (TRPV1) on primary afferent nociceptors[89] when signaling via their primary target is no longer available. Moreover, following tissue injury, CB_1_ blockade leads to decreased levels of pro-nociceptive cytokines, such as tumor necrosis factor-α (TNF-α)[90, 91] which could contribute to this phenomenon. Moreover, GAT358-induced blockade of nociception was occluded by GAT229, a CB_1_-PAM, indicating that a putative CB_1_ allosteric mechanism of action that mediates the effects of GAT358. GAT229 shows no independent activity at CB_1_ but potentiated the effect of several endogenous CB_1_ orthosteric agonists, consistent with a PAM profile[92, 93]. However, the effects of a CB_1_-PAM combined with a CB_1_-NAM on nociception have not been previously evaluated. As several studies have shown that the in vitro profile of other CB_1_ allosteric modulators may not necessarily translate in vivo the potential mechanism of action of such combinations must be parsimoniously interpreted [39, 94, 95]. Occlusion of the effects of GAT358 by a GAT229 may be the result of alternate mechanisms (e.g., physiological antagonism) that are not necessarily allosterically-mediated. More work is necessary to identify the exact mechanism or mechanisms involved.

Finally, physical dependence is a common side-effect of chronic opioid use and typically leads to a withdrawal syndrome following the cessation of drug use[96]. In our studies, GAT358 mitigated naloxone-precipitated opioid withdrawal but not putative spontaneous withdrawal in morphine-dependent mice. Our results do not preclude the possibility that GAT358 could produce a transient attenuation of spontaneous morphine withdrawal during temporal intervals that were not evaluated in the present study. Our results align with the observation that CB_1_KO mice show decreased severity of naloxone-precipitated morphine withdrawal[97, 98]. While chronic rimonabant treatment lessens the intensity of naloxone-precipitated opioid withdrawal[27, 28], it also produces CB_1_-dependent scratching behavior in naive[99–101] mice and opioid-like withdrawal effects in morphine-dependent animals[102]. Consequently, direct orthosteric CB_1_ blockade might produce adverse effects during withdrawal that further support the use of a CB_1_-NAM to overcome this unwanted side-effect of opioid use or rimonabant treatment.

Our results validate the use of a CB_1_-NAM to evaluate the hypothesis that CB_1_ blockade can facilitate acute antinociception without blocking opioid-mediated antinociception while suppressing other unwanted side-effects associated with chronic opioid use. Nonetheless, some potential caveats should be considered in interpreting our data. First, although GAT358 is a beta-arrestin-biased NAM, more work is necessary to determine if arrestin bias is necessary for the observed in vivo profile, particularly for antinociception. Second, because we evaluated the clinically relevant systemic route of drug administration, more work is necessary to determine the site of action of GAT358 by assessing the impact of intrathecal, local i.pl., and site-specific supraspinal injections. For example, the spinal cord dorsal horn is a key site in mediating the effects of CB_1_ receptor antagonists and a single intrathecal injection has been shown to produce antinociception in previous studies[30, 103]. Third, the antinociceptive effects of GAT358 in suppressing nociception induced by diverse inflammatory agents (capsaicin, complete Freund’s adjuvant (CFA) etc.), models of neuropathic pain (paclitaxel, traumatic nerve injury etc.) and with multiple modalities of cutaneous stimulation (mechanical, cold hypersensitivity etc.) need to be assessed. Finally, future studies will seek to extend these findings to female rodents, as the impact of sex on cannabinoids[104] and opioids[105–107] must be considered. Addressing these factors may increase the translational relevance of our findings.

In summary, our studies document that the CB_1_-NAM GAT358 reduces morphine antinociceptive tolerance, morphine-induced slowing of colonic motility, and somatic signs of naloxone-precipitated opioid withdrawal. Moreover, GAT358 produces anti-nociceptive effects in a formalin model of inflammatory pain in the presence and absence of opioids and also reduces the number of formalin-evoked Fos expressing cells in the lumbar spinal dorsal horn laminae associated with nociceptive processing by itself. We postulate that functionally selective CB_1_-NAMs such as GAT358 could potentially suppress aberrant signaling cascades involved in opioid-mediated side-effects, while preserving mechanisms essential for therapeutic analgesic effects.

Our results underscore the clinical potential of cannabinoid-opioid multimodal therapies and may provide a potentially novel approach for clinical pain management.

## Declaration of Interests

## Acknowledgements

This work is supported by the National Institutes of Health National Institute on Drug Abuse (NIDA) Grants DA047858 and DA041229 (A.G.H.), DA042584 (A.G.H.), DA027113 and EY024717 (G.A.T.), an Indiana Addiction Grand Challenge Grant (A.G.H.), a Gill Graduate Research Fellowship and the Harlan Scholars Research Program (V.I.).

## Disclosures

G.A.T. holds a patent on allosteric modulators of CB_1_ cannabinoid receptors (US9926275B2). V.I. is currently an employee of EMD Serono Inc. A.G. H. is a consultant for Anagin, Inc. None of the other authors report any conflicts of interest.

## CRediT author statement

**Vishakh Iyer:** Conceptualization, Methodology, Formal analysis, Investigation, Writing - Original Draft, Writing - Review & Editing, Visualization

**Shahin A. Saberi:** Conceptualization, Methodology, Formal analysis, Investigation, Visualization

**Romario Pacheco:** Conceptualization, Methodology, Formal analysis, Investigation, Visualization

**Emily Fender Sizemore:** Conceptualization, Methodology, Formal analysis, Investigation, Writing - Review & Editing, Visualization

**Sarah Stockman:** Formal analysis, Investigation, Visualization

**Abhijit Kulkarni:** Methodology, Formal analysis, Resources

**Lucas Cantwell:** Methodology, Formal analysis, Resources

**Ganesh A. Thakur:** Conceptualization, Methodology, Funding acquisition, Resources, Supervision, Writing - review & editing

**Andrea G. Hohmann:** Conceptualization, Methodology, Formal analysis, Visualization, Resources, Writing - Review & Editing, Supervision, Project administration, Funding acquisition

---

**Supplementary Fig. 1.**
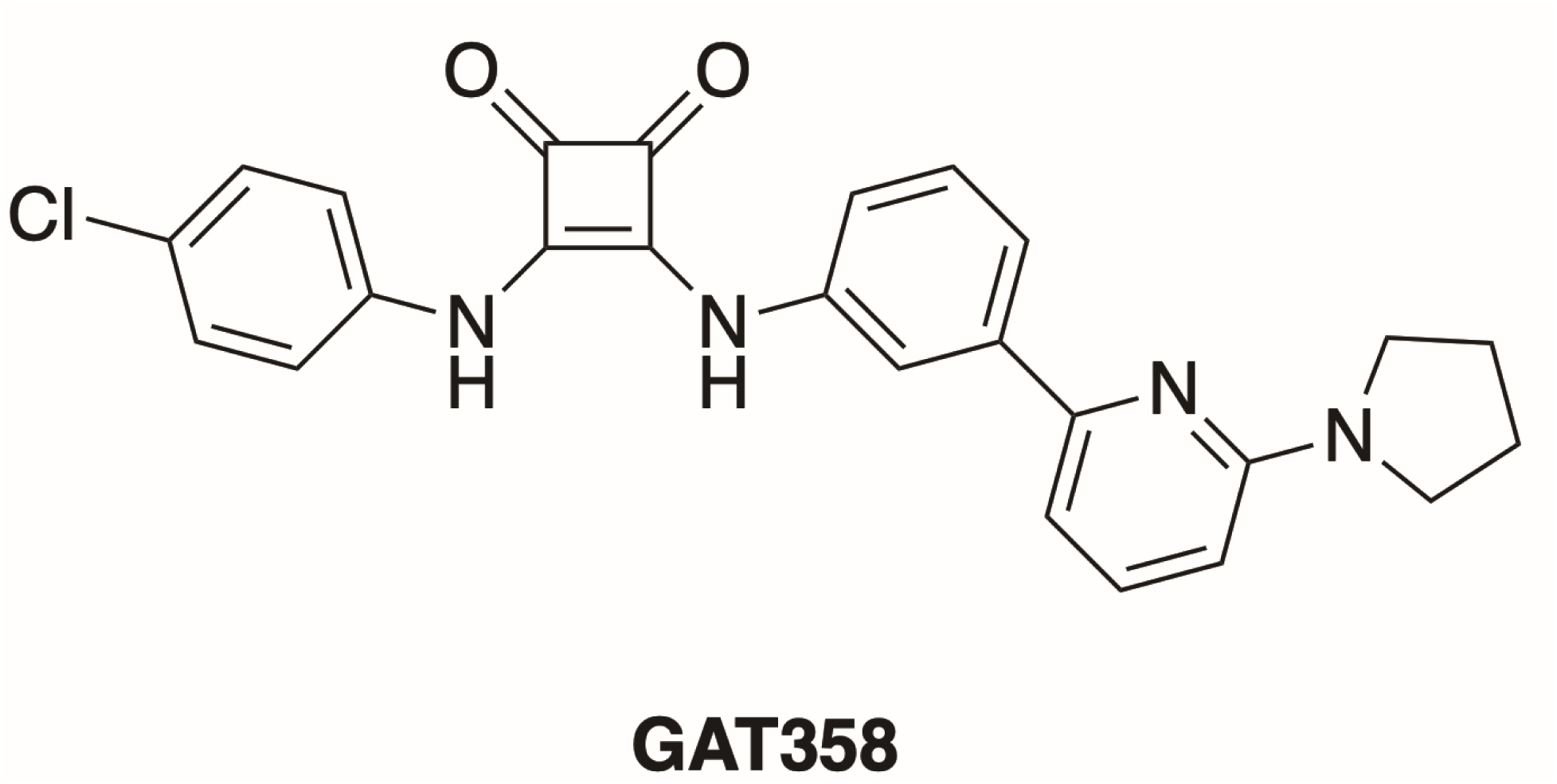
Chemical structure of GAT358. The figure shows the structure of the CB_1_-NAM, GAT358. Figures based on [45, 64, 92].

**Supplementary Fig. 2.**
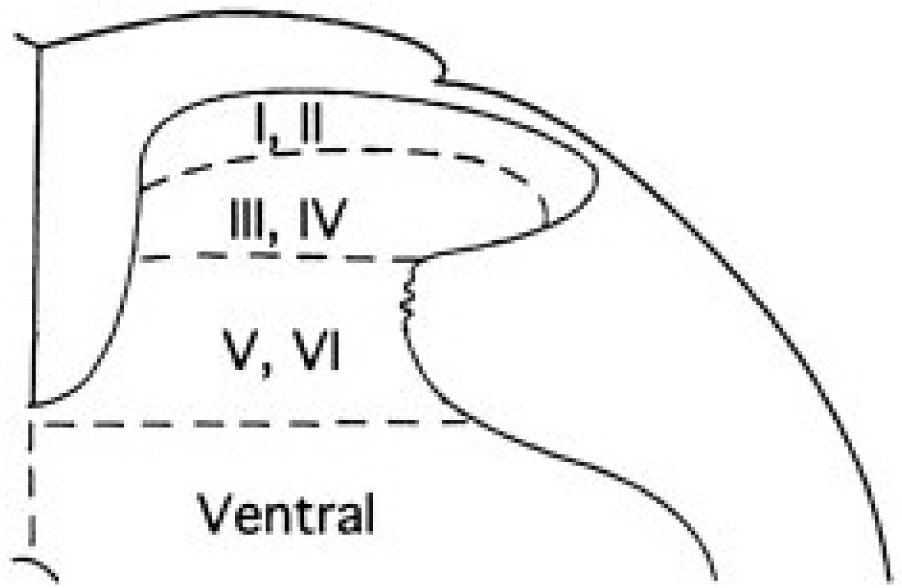
Diagram of a lumbar spinal cord hemi-section showing laminar subdivisions used to quantify formalin-evoked Fos protein expression. The figure shows laminae I, II (superficial dorsal horn), laminae III, IV (nucleus proprius), laminae V,VI (neck region of the dorsal horn) and the ventral horn of the lumbar spinal cord used to quantify formalin-evoked Fos protein expression (adapted from [108, 109]).

**Supplementary Fig. 3.**
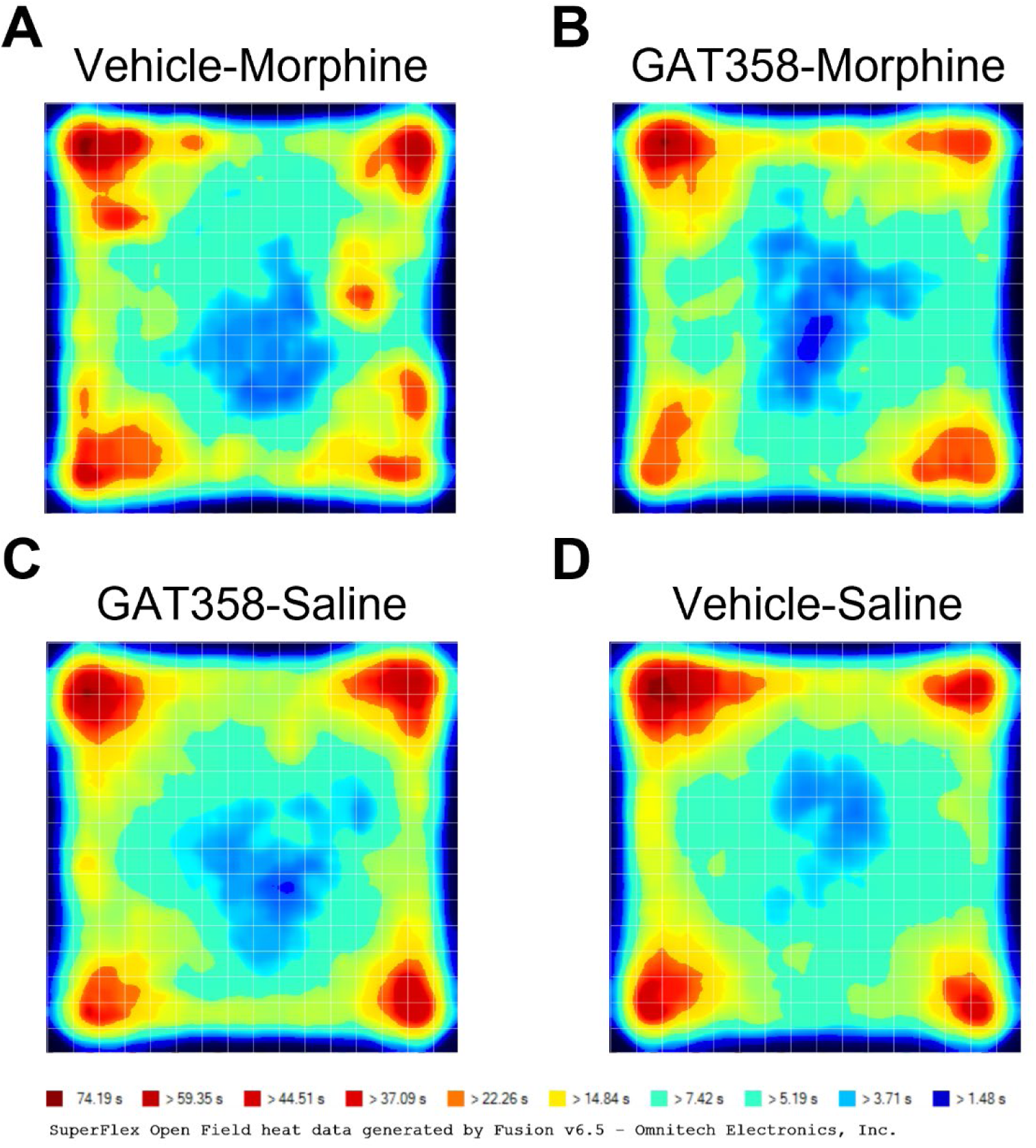
Cumulative heat maps of locomotor activity in morphine-dependent mice undergoing spontaneous withdrawal. Cumulative heat maps of locomotor activity in morphine-dependent mice evaluated in the activity meter to assess possible spontaneous withdrawal. Behavior was quantified over a 30-min observation interval 24 h after the last i.p. injection of **(A)** Vehicle + Morphine-, **(B)** GAT358 + Morphine-, **(C)** GAT358 + Saline-, and **(D)** Vehicle + Saline-treated mice. Colors are indicative of the time spent in different zones of the activity meter.

## References

[1] T.J. Cicero, M.S. Ellis, The prescription opioid epidemic: a review of qualitative studies on the progression from initial use to abuse, Dialogues Clin Neurosci 19(3) (2017) 259–269.

[2] F. Ahmad, L. Rossen, P. Sutton, Provisional drug overdose death counts National Center for Health Statistics. 2021, Statistics, Centers for Disease Control and Prevention (CDC). (2021).

[3] N. Wilson, M. Kariisa, P. Seth, H.t. Smith, N.L. Davis, Drug and Opioid-Involved Overdose Deaths -United States, 2017-2018, MMWR Morb Mortal Wkly Rep 69(11) (2020) 290–297.

[4] R.J. Wickham, Cancer Pain Management: Opioid Analgesics, Part 2, J Adv Pract Oncol 8(6) (2017) 588-607.

[5] M. Shkodra, A. Caraceni, Treatment of Neuropathic Pain Directly Due to Cancer: An Update, Cancers (Basel) 14(8) (2022).

[6] N.B. Finnerup, N. Attal, S. Haroutounian, E. McNicol, R. Baron, R.H. Dworkin, I. Gilron, M. Haanpaa, P. Hansson, T.S. Jensen, P.R. Kamerman, K. Lund, A. Moore, S.N. Raja, A.S. Rice, M. Rowbotham, E. Sena, P. Siddall, B.H. Smith, M. Wallace, Pharmacotherapy for neuropathic pain in adults: a systematic review and meta-analysis, Lancet Neurol 14(2) (2015) 162–73.

[7] R.N. Jamison, J. Mao, Opioid Analgesics, Mayo Clinic proceedings 90(7) (2015) 957–68.

[8] H.L. Fields, The doctor’s dilemma: opiate analgesics and chronic pain, Neuron 69(4) (2011) 591–4.

[9] A.D. Furlan, J.A. Sandoval, A. Mailis-Gagnon, E. Tunks, Opioids for chronic noncancer pain: a meta-analysis of effectiveness and side effects, CMAJ 174(11) (2006) 1589–94.

[10] W.M. Compton, M. Boyle, E. Wargo, Prescription opioid abuse: Problems and responses, Prev Med 80 (2015) 5–9.

[11] P. Skolnick, The Opioid Epidemic: Crisis and Solutions, Annu Rev Pharmacol Toxicol 58 (2018) 143–159.

[12] C.A. Zavala, A.C. Thomaz, V. Iyer, K. Mackie, A.G. Hohmann, Cannabinoid CB(2) Receptor Activation Attenuates Fentanyl-Induced Respiratory Depression, Cannabis Cannabinoid Res 6(5) (2021) 389–400.

[13] R.G. Pertwee, Cannabinoid receptors and pain, Prog Neurobiol 63(5) (2001) 569–611.

[14] B.E. Perron, K. Bohnert, A.K. Perone, M.O. Bonn-Miller, M. Ilgen, Use of prescription pain medications among medical cannabis patients: comparisons of pain levels, functioning, and patterns of alcohol and other drug use, J Stud Alcohol Drugs 76(3) (2015) 406–13.

[15] T. Yamaguchi, Y. Hagiwara, H. Tanaka, T. Sugiura, K. Waku, Y. Shoyama, S. Watanabe, T. Yamamoto, Endogenous cannabinoid, 2-arachidonoylglycerol, attenuates naloxone-precipitated withdrawal signs in morphine-dependent mice, Brain Res 909(1-2) (2001) 121–6.

[16] D.L. Cichewicz, S.P. Welch, Modulation of oral morphine antinociceptive tolerance and naloxone-precipitated withdrawal signs by oral Delta 9-tetrahydrocannabinol, J Pharmacol Exp Ther 305(3) (2003) 812–7.

[17] J.L. Scavone, R.C. Sterling, E.J. Van Bockstaele, Cannabinoid and opioid interactions: implications for opiate dependence and withdrawal, Neuroscience 248 (2013) 637–54.

[18] U. Taschler, T.O. Eichmann, F.P. Radner, G.F. Grabner, H. Wolinski, M. Storr, A. Lass, R. Schicho, R. Zimmermann, Monoglyceride lipase deficiency causes desensitization of intestinal cannabinoid receptor type 1 and increased colonic mu-opioid receptor sensitivity, Br J Pharmacol 172(17) (2015) 4419–29.

[19] A.R. Wilson, L. Maher, M.M. Morgan, Repeated cannabinoid injections into the rat periaqueductal gray enhance subsequent morphine antinociception, Neuropharmacology 55(7) (2008) 1219–25.

[20] R.A. Slivicki, V. Iyer, S.S. Mali, S. Garai, G.A. Thakur, J.D. Crystal, A.G. Hohmann, Positive Allosteric Modulation of CB(1) Cannabinoid Receptor Signaling Enhances Morphine Antinociception and Attenuates Morphine Tolerance Without Enhancing Morphine-Induced Dependence or Reward, Front Mol Neurosci 13 (2020) 54.

[21] M.L. Cox, V.L. Haller, S.P. Welch, Synergy between delta9-tetrahydrocannabinol and morphine in the arthritic rat, Eur J Pharmacol 567(1-2) (2007) 125–30.

[22] J.L. Wilkerson, S. Ghosh, M. Mustafa, R.A. Abdullah, M.J. Niphakis, R. Cabrera, R.L. Maldonado, B.F. Cravatt, A.H. Lichtman, The endocannabinoid hydrolysis inhibitor SA-57: Intrinsic antinociceptive effects, augmented morphine-induced antinociception, and attenuated heroin seeking behavior in mice, Neuropharmacology 114 (2017) 156–167.

[23] J.L. Wilkerson, M.J. Niphakis, T.W. Grim, M.A. Mustafa, R.A. Abdullah, J.L. Poklis, W.L. Dewey, H. Akbarali, M.L. Banks, L.E. Wise, B.F. Cravatt, A.H. Lichtman, The Selective Monoacylglycerol Lipase Inhibitor MJN110 Produces Opioid-Sparing Effects in a Mouse Neuropathic Pain Model, J Pharmacol Exp Ther 357(1) (2016) 145–56.

[24] K.L. Wills, L.A. Parker, Effect of Pharmacological Modulation of the Endocannabinoid System on Opiate Withdrawal: A Review of the Preclinical Animal Literature, Front Pharmacol 7 (2016) 187.

[25] K. Befort, Interactions of the opioid and cannabinoid systems in reward: Insights from knockout studies, Front Pharmacol 6 (2015) 6.

[26] T. Trang, M. Sutak, K. Jhamandas, Involvement of cannabinoid (CB1)-receptors in the development and maintenance of opioid tolerance, Neuroscience 146(3) (2007) 1275–88.

[27] T. Rubino, P. Massi, D. Vigano, D. Fuzio, D. Parolaro, Long-term treatment with SR141716A, the CB1 receptor antagonist, influences morphine withdrawal syndrome, Life Sci 66(22) (2000) 2213–9.

[28] M. Mas-Nieto, B. Pommier, E.T. Tzavara, A. Caneparo, S. Da Nascimento, G. Le Fur, B.P. Roques, F. Noble, Reduction of opioid dependence by the CB(1) antagonist SR141716A in mice: evaluation of the interest in pharmacotherapy of opioid addiction, Br J Pharmacol 132(8) (2001) 1809–16.

[29] T. Croci, E. Zarini, Effect of the cannabinoid CB1 receptor antagonist rimonabant on nociceptive responses and adjuvant-induced arthritis in obese and lean rats, Br J Pharmacol 150(5) (2007) 559–66.

[30] A.J. Pernia-Andrade, A. Kato, R. Witschi, R. Nyilas, I. Katona, T.F. Freund, M. Watanabe, J. Filitz, W. Koppert, J. Schuttler, G. Ji, V. Neugebauer, G. Marsicano, B. Lutz, H. Vanegas, H.U. Zeilhofer, Spinal endocannabinoids and CB1 receptors mediate C-fiber-induced heterosynaptic pain sensitization, Science 325(5941) (2009) 760–4.

[31] P. Beaulieu, T. Bisogno, S. Punwar, W.P. Farquhar-Smith, G. Ambrosino, V. Di Marzo, A.S. Rice, Role of the endogenous cannabinoid system in the formalin test of persistent pain in the rat, Eur J Pharmacol 396(2-3) (2000) 85–92.

[32] M. Ueda, H. Iwasaki, S. Wang, E. Murata, K.Y. Poon, J. Mao, J.A. Martyn, Cannabinoid receptor type 1 antagonist, AM251, attenuates mechanical allodynia and thermal hyperalgesia after burn injury, Anesthesiology 121(6) (2014) 1311–9.

[33] E. Kirilly, X. Gonda, G. Bagdy, CB1 receptor antagonists: new discoveries leading to new perspectives, Acta Physiol (Oxf) 205(1) (2012) 41–60.

[34] D. Jones, End of the line for cannabinoid receptor 1 as an anti-obesity target?, Nat Rev Drug Discov 7(12) (2008) 961–2.

[35] F.A. Moreira, M. Grieb, B. Lutz, Central side-effects of therapies based on CB1 cannabinoid receptor agonists and antagonists: focus on anxiety and depression, Best Pract Res Clin Endocrinol Metab 23(1) (2009) 133–44.

[36] K. Johansson, K. Neovius, S. DeSantis, S. Rössner, M. Neovius, Discontinuation due to adverse events in randomized trials of orlistat, sibutramine and rimonabant: a meta-analysis, obesity reviews 10(5) (2009) 564–575.

[37] M.R. Price, G.L. Baillie, A. Thomas, L.A. Stevenson, M. Easson, R. Goodwin, A. McLean, L. McIntosh, G. Goodwin, G. Walker, P. Westwood, J. Marrs, F. Thomson, P. Cowley, A. Christopoulos, R.G. Pertwee, R.A. Ross, Allosteric modulation of the cannabinoid CB1 receptor, Mol Pharmacol 68(5) (2005) 1484–95.

[38] D.M. Shore, G.L. Baillie, D.H. Hurst, F. Navas, 3rd, H.H. Seltzman, J.P. Marcu, M.E. Abood, R.A. Ross, P.H. Reggio, Allosteric modulation of a cannabinoid G protein-coupled receptor: binding site elucidation and relationship to G protein signaling, J Biol Chem 289(9) (2014) 5828–45.

[39] T.F. Gamage, B.M. Ignatowska-Jankowska, J.L. Wiley, M. Abdelrahman, L. Trembleau, I.R. Greig, G.A. Thakur, R. Tichkule, J. Poklis, R.A. Ross, R.G. Pertwee, A.H. Lichtman, In-vivo pharmacological evaluation of the CB1-receptor allosteric modulator Org-27569, Behav Pharmacol 25(2) (2014) 182–5.

[40] L. Jing, Y. Qiu, Y. Zhang, J.X. Li, Effects of the cannabinoid CB(1) receptor allosteric modulator ORG 27569 on reinstatement of cocaine- and methamphetamine-seeking behavior in rats, Drug Alcohol Depend 143 (2014) 251–6.

[41] J.G. Horswill, U. Bali, S. Shaaban, J.F. Keily, P. Jeevaratnam, A.J. Babbs, C. Reynet, P. Wong Kai In, PSNCBAM-1, a novel allosteric antagonist at cannabinoid CB1 receptors with hypophagic effects in rats, Br J Pharmacol 152(5) (2007) 805–14.

[42] N.T. Burford, J.R. Traynor, A. Alt, Positive allosteric modulators of the mu-opioid receptor: a novel approach for future pain medications, Br J Pharmacol 172(2) (2015) 277–86.

[43] L. Khurana, K. Mackie, D. Piomelli, D.A. Kendall, Modulation of CB1 cannabinoid receptor by allosteric ligands: Pharmacology and therapeutic opportunities, Neuropharmacology 124 (2017) 3–12.

[44] P.L. Raux, G. Drutel, J.M. Revest, M. Vallee, New perspectives on the role of the neurosteroid pregnenolone as an endogenous regulator of type-1 cannabinoid receptor (CB1R) activity and function, J Neuroendocrinol (2021) e13034.

[45] V. Iyer, C. Rangel-Barajas, T.J. Woodward, A. Kulkarni, L. Cantwell, J.D. Crystal, K. Mackie, G.V. Rebec, G.A. Thakur, A.G. Hohmann, Negative allosteric modulation of CB(1) cannabinoid receptor signaling suppresses opioid-mediated reward, Pharmacol Res 185 (2022) 106474.

[46] M. Zimmermann, Ethical guidelines for investigations of experimental pain in conscious animals, Pain 16(2) (1983) 109–10.

[47] N. Percie du Sert, A. Ahluwalia, S. Alam, M.T. Avey, M. Baker, W.J. Browne, A. Clark, I.C. Cuthill, U. Dirnagl, M. Emerson, P. Garner, S.T. Holgate, D.W. Howells, V. Hurst, N.A. Karp, S.E. Lazic, K. Lidster, C.J. MacCallum, M. Macleod, E.J. Pearl, O.H. Petersen, F. Rawle, P. Reynolds, K. Rooney, E.S. Sena, S.D. Silberberg, T. Steckler, H. Wurbel, Reporting animal research: Explanation and elaboration for the ARRIVE guidelines 2.0, PLoS Biol 18(7) (2020) e3000411.

[48] L.M. Console-Bram, P. Zhao, M.E. Abood, Protocols and Good Operating Practices in the Study of Cannabinoid Receptors, Methods Enzymol 593 (2017) 23–42.

[49] R.A. Slivicki, V. Iyer, S.S. Mali, S. Garai, G.A. Thakur, J.D. Crystal, A.G. Hohmann, Positive Allosteric Modulation of CB1 Cannabinoid Receptor Signaling Enhances Morphine Antinociception and Attenuates Morphine Tolerance Without Enhancing Morphine-Induced Dependence or Reward, Frontiers in Molecular Neuroscience 13(54) (2020).

[50] R.A. Slivicki, S.A. Saberi, V. Iyer, V.K. Vemuri, A. Makriyannis, A.G. Hohmann, Brain-Permeant and - Impermeant Inhibitors of Fatty Acid Amide Hydrolase Synergize with the Opioid Analgesic Morphine to Suppress Chemotherapy-Induced Neuropathic Nociception Without Enhancing Effects of Morphine on Gastrointestinal Transit, J Pharmacol Exp Ther 367(3) (2018) 551–563.

[51] V. Iyer, R.A. Slivicki, A.C. Thomaz, J.D. Crystal, K. Mackie, A.G. Hohmann, The cannabinoid CB2 receptor agonist LY2828360 synergizes with morphine to suppress neuropathic nociception and attenuates morphine reward and physical dependence, Eur J Pharmacol 886 (2020) 173544.

[52] I. Oliva, S.A. Saberi, C. Rangel-Barajas, V. Iyer, K.D. Bunner, Y.Y. Lai, P.M. Kulkarni, S. Garai, G.A. Thakur, J.D. Crystal, G.V. Rebec, A.G. Hohmann, Inhibition of PSD95-nNOS protein-protein interactions decreases morphine reward and relapse vulnerability in rats, Addict Biol 27(5) (2022) e13220.

[53] V. Iyer, T.J. Woodward, R. Pacheco, A.G. Hohmann, A limited access oral oxycodone paradigm produces physical dependence and mesocorticolimbic region-dependent increases in DeltaFosB expression without preference, Neuropharmacology 205 (2022) 108925.

[54] L. Liu, J.K. Coller, L.R. Watkins, A.A. Somogyi, M.R. Hutchinson, Naloxone-precipitated morphine withdrawal behavior and brain IL-1beta expression: comparison of different mouse strains, Brain Behav Immun 25(6) (2011) 1223–32.

[55] L.M. Carey, W.H. Lee, T. Gutierrez, P.M. Kulkarni, G.A. Thakur, Y.Y. Lai, A.G. Hohmann, Small molecule inhibitors of PSD95-nNOS protein-protein interactions suppress formalin-evoked Fos protein expression and nociceptive behavior in rats, Neuroscience 349 (2017) 303–317.

[56] W.H. Lee, L.M. Carey, L.L. Li, Z. Xu, Y.Y. Lai, M.J. Courtney, A.G. Hohmann, ZLc002, a putative small-molecule inhibitor of nNOS interaction with NOS1AP, suppresses inflammatory nociception and chemotherapy-induced neuropathic pain and synergizes with paclitaxel to reduce tumor cell viability, Mol Pain 14 (2018) 1744806918801224.

[57] A.C. Thomaz, V. Iyer, T.J. Woodward, A.G. Hohmann, Fecal microbiota transplantation and antibiotic treatment attenuate naloxone-precipitated opioid withdrawal in morphine-dependent mice, Exp Neurol 343 (2021) 113787.

[58] R.A. Slivicki, V. Iyer, S.S. Mali, S. Garai, G.A. Thakur, J.D. Crystal, A.G. Hohmann, Positive Allosteric Modulation of CB1 Cannabinoid Receptor Signaling Enhances Morphine Antinociception and Attenuates Morphine Tolerance Without Enhancing Morphine-Induced Dependence or Reward, Front Mol Neurosci 13 (2020) 54.

[59] O. Friard, M. Gamba, BORIS: a free, versatile open-source event-logging software for video/audio coding and live observations, Methods in Ecology and Evolution 7(11) (2016) 1325–1330.

[60] D.S. McDevitt, G. McKendrick, N.M. Graziane, Anterior cingulate cortex is necessary for spontaneous opioid withdrawal and withdrawal-induced hyperalgesia in male mice, Neuropsychopharmacology 46(11) (2021) 1990–1999.

[61] F. Papaleo, A. Contarino, Gender- and morphine dose-linked expression of spontaneous somatic opiate withdrawal in mice, Behav Brain Res 170(1) (2006) 110–8.

[62] L.M. Carey, R.A. Slivicki, E. Leishman, B. Cornett, K. Mackie, H. Bradshaw, A.G. Hohmann, A pro-nociceptive phenotype unmasked in mice lacking fatty-acid amide hydrolase, Mol Pain 12 (2016).

[63] K. Rasmussen, D.A. White, J.B. Acri, NIDA’s medication development priorities in response to the Opioid Crisis: ten most wanted, Neuropsychopharmacology 44(4) (2019) 657–659.

[64] P. Morales, P. Goya, N. Jagerovic, L. Hernandez-Folgado, Allosteric Modulators of the CB1 Cannabinoid Receptor: A Structural Update Review, Cannabis Cannabinoid Res 1(1) (2016) 22–30.

[65] G.A. Thakur, R.B. Tichkule, P.M. Kulkarni, A.R. Kulkarni, Allosteric modulators of the cannabinoid 1 receptor, U.S. Patent US9926275B2, Northeastern University, U.S. Patent Office, 2018.

[66] M. Pappagallo, Incidence, prevalence, and management of opioid bowel dysfunction, Am J Surg 182(5A Suppl) (2001) 11S–18S.

[67] Y.S. Choi, J.A. Billings, Opioid antagonists: a review of their role in palliative care, focusing on use in opioid-related constipation, J Pain Symptom Manage 24(1) (2002) 71–90.

[68] T.J. Bell, S.J. Panchal, C. Miaskowski, S.C. Bolge, T. Milanova, R. Williamson, The prevalence, severity, and impact of opioid-induced bowel dysfunction: results of a US and European Patient Survey (PROBE 1), Pain Med 10(1) (2009) 35–42.

[69] G. Aviello, B. Romano, A.A. Izzo, Cannabinoids and gastrointestinal motility: animal and human studies, Eur Rev Med Pharmacol Sci 12 Suppl 1 (2008) 81–93.

[70] R.G. Pertwee, Cannabinoids and the gastrointestinal tract, Gut 48(6) (2001) 859–67.

[71] A.A. Izzo, L. Pinto, F. Borrelli, R. Capasso, N. Mascolo, F. Capasso, Central and peripheral cannabinoid modulation of gastrointestinal transit in physiological states or during the diarrhoea induced by croton oil, Br J Pharmacol 129(8) (2000) 1627–32.

[72] G. Colombo, R. Agabio, C. Lobina, R. Reali, G.L. Gessa, Cannabinoid modulation of intestinal propulsion in mice, Eur J Pharmacol 344(1) (1998) 67–9.

[73] M.A. Carai, G. Colombo, G.L. Gessa, R. Yalamanchili, B.S. Basavarajappa, B.L. Hungund, Investigation on the relationship between cannabinoid CB1 and opioid receptors in gastrointestinal motility in mice, Br J Pharmacol 148(8) (2006) 1043–50.

[74] E.O. Dumas, G.M. Pollack, Opioid tolerance development: a pharmacokinetic/pharmacodynamic perspective, AAPS J 10(4) (2008) 537–51.

[75] A. Altun, E. Ozdemir, K. Yildirim, S. Gursoy, N. Durmus, I. Bagcivan, The effects of endocannabinoid receptor agonist anandamide and antagonist rimonabant on opioid analgesia and tolerance in rats, Gen Physiol Biophys 34(4) (2015) 433–40.

[76] A. Altun, K. Yildirim, E. Ozdemir, I. Bagcivan, S. Gursoy, N. Durmus, Attenuation of morphine antinociceptive tolerance by cannabinoid CB1 and CB2 receptor antagonists, J Physiol Sci 65(5) (2015) 407–15.

[77] G. Roussy, M.A. Dansereau, L. Dore-Savard, K. Belleville, N. Beaudet, E. Richelson, P. Sarret, Spinal NTS1 receptors regulate nociceptive signaling in a rat formalin tonic pain model, J Neurochem 105(4) (2008) 1100–14.

[78] S. Puig, L.S. Sorkin, Formalin-evoked activity in identified primary afferent fibers: systemic lidocaine suppresses phase-2 activity, Pain 64(2) (1996) 345–355.

[79] J.L.K. Hylden, R.L. Nahin, R.J. Traub, R. Dubner, Expansion of receptive fields of spinal lamina I projection neurons in rats with unilateral adjuvant-induced inflammation: the contribution of dorsal horn mechanisms, Pain 37(2) (1989) 229–243.

[80] P. Lebrun, J. Manil, F. Colin, Formalin-induced central sensitization in the rat: somatosensory evoked potential data, Neurosci Lett 283(2) (2000) 113–6.

[81] K.B. Franklin, F.V. Abbott, Pentobarbital, diazepam, and ethanol abolish the interphase diminution of pain in the formalin test: evidence for pain modulation by GABAA receptors, Pharmacol Biochem Behav 46(3) (1993) 661–6.

[82] M. Kaneko, D.L. Hammond, Role of spinal gamma-aminobutyric acidA receptors in formalin-induced nociception in the rat, J Pharmacol Exp Ther 282(2) (1997) 928–38.

[83] D. Menetrey, A. Gannon, J.D. Levine, A.I. Basbaum, Expression of c-fos protein in interneurons and projection neurons of the rat spinal cord in response to noxious somatic, articular, and visceral stimulation, J Comp Neurol 285(2) (1989) 177–95.

[84] W.P. Farquhar-Smith, M. Egertova, E.J. Bradbury, S.B. McMahon, A.S. Rice, M.R. Elphick, Cannabinoid CB(1) receptor expression in rat spinal cord, Mol Cell Neurosci 15(6) (2000) 510–21.

[85] A.G. Hohmann, E.M. Briley, M. Herkenham, Pre- and postsynaptic distribution of cannabinoid and mu opioid receptors in rat spinal cord, Brain Res 822(1-2) (1999) 17–25.

[86] A.G. Hohmann, M. Herkenham, Regulation of cannabinoid and mu opioid receptors in rat lumbar spinal cord following neonatal capsaicin treatment, Neurosci Lett 252(1) (1998) 13–6.

[87] R. Nyilas, L.C. Gregg, K. Mackie, M. Watanabe, A. Zimmer, A.G. Hohmann, I. Katona, Molecular architecture of endocannabinoid signaling at nociceptive synapses mediating analgesia, Eur J Neurosci 29(10) (2009) 1964–78.

[88] J.M. Walker, S.M. Huang, N.M. Strangman, K. Tsou, M.C. Sanudo-Pena, Pain modulation by release of the endogenous cannabinoid anandamide, Proc Natl Acad Sci U S A 96(21) (1999) 12198–203.

[89] B. Costa, Rimonabant: more than an anti-obesity drug?, Br J Pharmacol 150(5) (2007) 535–7.

[90] B. Costa, A.E. Trovato, M. Colleoni, G. Giagnoni, E. Zarini, T. Croci, Effect of the cannabinoid CB1 receptor antagonist, SR141716, on nociceptive response and nerve demyelination in rodents with chronic constriction injury of the sciatic nerve, Pain 116(1-2) (2005) 52–61.

[91] T. Croci, M. Landi, A.M. Galzin, P. Marini, Role of cannabinoid CB1 receptors and tumor necrosis factor-alpha in the gut and systemic anti-inflammatory activity of SR 141716 (rimonabant) in rodents, Br J Pharmacol 140(1) (2003) 115–22.

[92] R.B. Laprairie, P.M. Kulkarni, J.R. Deschamps, M.E.M. Kelly, D.R. Janero, M.G. Cascio, L.A. Stevenson, R.G. Pertwee, T.P. Kenakin, E.M. Denovan-Wright, G.A. Thakur, Enantiospecific Allosteric Modulation of Cannabinoid 1 Receptor, ACS Chem Neurosci 8(6) (2017) 1188–1203.

[93] D. Thapa, E.A. Cairns, A.M. Szczesniak, P.M. Kulkarni, A.J. Straiker, G.A. Thakur, M.E.M. Kelly, Allosteric Cannabinoid Receptor 1 (CB1) Ligands Reduce Ocular Pain and Inflammation, Molecules 25(2) (2020).

[94] T.F. Gamage, C.E. Farquhar, T.W. Lefever, B.F. Thomas, T. Nguyen, Y. Zhang, J.L. Wiley, The great divide: Separation between in vitro and in vivo effects of PSNCBAM-based CB1 receptor allosteric modulators, Neuropharmacology 125 (2017) 365–375.

[95] D. Lu, S.S. Immadi, Z. Wu, D.A. Kendall, Translational potential of allosteric modulators targeting the cannabinoid CB1 receptor, Acta Pharmacol Sin 40(3) (2019) 324–335.

[96] G.F. Koob, R. Maldonado, L. Stinus, Neural substrates of opiate withdrawal, Trends Neurosci 15(5) (1992) 186–91.

[97] A.H. Lichtman, S.M. Sheikh, H.H. Loh, B.R. Martin, Opioid and cannabinoid modulation of precipitated withdrawal in delta(9)-tetrahydrocannabinol and morphine-dependent mice, J Pharmacol Exp Ther 298(3) (2001) 1007–14.

[98] C. Ledent, O. Valverde, G. Cossu, F. Petitet, J.F. Aubert, F. Beslot, G.A. Bohme, A. Imperato, T. Pedrazzini, B.P. Roques, G. Vassart, W. Fratta, M. Parmentier, Unresponsiveness to cannabinoids and reduced addictive effects of opiates in CB1 receptor knockout mice, Science 283(5400) (1999) 401–4.

[99] F. Rodriguez de Fonseca, M.R. Carrera, M. Navarro, G.F. Koob, F. Weiss, Activation of corticotropin-releasing factor in the limbic system during cannabinoid withdrawal, Science 276(5321) (1997) 2050–4.

[100] M.D. Aceto, S.M. Scates, J.A. Lowe, B.R. Martin, Cannabinoid precipitated withdrawal by the selective cannabinoid receptor antagonist, SR 141716A, Eur J Pharmacol 282(1-3) (1995) R1–2.

[101] M.D. Aceto, S.M. Scates, J.A. Lowe, B.R. Martin, Dependence on delta 9-tetrahydrocannabinol: studies on precipitated and abrupt withdrawal, J Pharmacol Exp Ther 278(3) (1996) 1290–5.

[102] M. Navarro, J. Chowen, A.C.M. Rocio, I. del Arco, M.A. Villanua, Y. Martin, A.J. Roberts, G.F. Koob, F.R. de Fonseca, CB1 cannabinoid receptor antagonist-induced opiate withdrawal in morphine-dependent rats, Neuroreport 9(15) (1998) 3397–402.

[103] A.S. Heimann, I. Gomes, C.S. Dale, R.L. Pagano, A. Gupta, L.L. de Souza, A.D. Luchessi, L.M. Castro, R. Giorgi, V. Rioli, E.S. Ferro, L.A. Devi, Hemopressin is an inverse agonist of CB1 cannabinoid receptors, Proc Natl Acad Sci U S A 104(51) (2007) 20588–93.

[104] Z.D. Cooper, R.M. Craft, Sex-Dependent Effects of Cannabis and Cannabinoids: A Translational Perspective, Neuropsychopharmacology 43(1) (2018) 34–51.

[105] J.B. Becker, G.F. Koob, Sex Differences in Animal Models: Focus on Addiction, Pharmacol Rev 68(2) (2016) 242–63.

[106] S.A. Bobzean, A.K. DeNobrega, L.I. Perrotti, Sex differences in the neurobiology of drug addiction, Exp Neurol 259 (2014) 64–74.

[107] M. Serdarevic, C.W. Striley, L.B. Cottler, Sex differences in prescription opioid use, Curr Opin Psychiatry 30(4) (2017) 238–246.

[108] A.G. Nackley, R.L. Suplita, 2nd, A.G. Hohmann, A peripheral cannabinoid mechanism suppresses spinal fos protein expression and pain behavior in a rat model of inflammation, Neuroscience 117(3) (2003) 659–70.

[109] C. Watson, G. Paxinos, G. Kayalioglu, C. Heise, Chapter 16 - Atlas of the Mouse Spinal Cord, in: C. Watson, G. Paxinos, G. Kayalioglu (Eds.), The Spinal Cord, Academic Press, San Diego, 2009, pp. 308–379.

